# Dynein activation in vivo is regulated by the nucleotide states of its AAA3 domain

**DOI:** 10.1101/2021.04.12.439451

**Authors:** Rongde Qiu, Jun Zhang, Jeremy D. Rotty, Xin Xiang

## Abstract

Cytoplasmic dynein is activated by dynactin and cargo adapters in vitro, and the activation also needs LIS1 (Lissencephaly 1) in vivo. How this process is regulated remains unclear. Here we found in *Aspergillus nidulans* that a dynein AAA4 arginine-finger mutation bypasses the requirement of LIS1 for dynein activation driven by the early endosomal adapter HookA. As the AAA4 arginine-finger is implicated in AAA3 ATP hydrolysis, we examined AAA3 mutants defective in ATP binding and hydrolysis respectively. Astonishingly, blocking AAA3 ATP hydrolysis allows dynein activation by dynactin in the absence of LIS1 or HookA. As a consequence, dynein accumulates at microtubule minus ends while early endosomes stay near the plus ends. On the other hand, blocking AAA3 ATP binding abnormally prevents LIS1 from being dissociated from dynein upon motor activation. Thus, the AAA3 ATPase cycle regulates the coordination between dynein activation and cargo binding as well as the dynamic dynein-LIS1 interaction.

## Introduction

Cytoplasmic dynein-1 (called “dynein” hereafter for simplicity) is a minus-end-directed microtubule motor that transports a variety of cellular cargoes as well as viruses, and dysfunction of dynein or its regulators is linked to neurodevelopmental disorders and neurogenerative diseases [1-6]. Dynein is a multi-protein complex containing heavy chains, intermediate chains, light intermediate chains and light chains [7]. The two dynein heavy chains form a homodimer, and each heavy chain contains a motor ring with six AAA+ (ATPases Associated with diverse cellular Activities) domains, a stalk connecting the microtubule-binding domain and a buttress that interacting with the stalk [8-19]. At the N-terminus of AAA1 is the linker domain, whose conformational change is involved in dynein’s mechanochemical cycle [13, 17, 20-22], and at the N-terminus of the linker is the dynein tail that binds other dynein subunits and the multi-component dynactin complex [7, 23-32]. The dynactin complex and specific cargo adapters allow dynein to associate with cargoes such as early endosomes [2, 33-41]. Importantly, several coiled-coils-containing cargo adapters, including the BicD family and the Hook family of proteins, activate the processive movement of mammalian dynein in the presence of dynactin [24, 42-44]. Dynactin and cargo adapter-mediated dynein activation is caused by a series of conformational changes [32]. The dynein dimer initially forms an autoinhibited “phi” conformation [32, 45, 46], which can be switched to an open conformation with the motor domains being separated from each other; Binding of dynactin and a cargo adapter to the dynein tails turns the dynein dimer to a parallel configuration, allowing it to walk along a microtubule [32]. Cargo adapter-mediated dynein activation is a useful mechanism that allows dynein to be activated only after it binds its cargo. Function of the cargo adapter has been implicated in enhancing the interaction between purified mammalian dynein and dynactin [24, 42, 43, 47]. However, since dynein and dynactin interact before cargo binding in some in vivo systems [48, 49], additional regulatory mechanisms may exist to ensure that dynein is not activated by dynactin alone before cargo binding.

The minus-end-directed movement of dynein requires multiple elements of the dynein heavy chain, including the AAA1-6 domains, the linker, the buttress, the stalk and its distal microtubule-binding domain [20, 50-53]. Although AAA1-4 domains can all bind ATP [54], only AAA1, 3, and 4 hydrolyze ATP, with AAA1 ATP hydrolysis being the most critical for driving motility [15, 17, 20, 55]. ATP binding at AAA1 drives the conformational change of the AAA ring and makes the ring closer [11, 22], and it also causes the linker to change from an extended conformation with its N-terminus docked at AAA5 to a bent conformation with its N-terminus moved to AAA2/3 [17, 22], and these changes are transmitted to buttress and stalk, causing the stalk to shift its coiled-coil registry, which switches the microtubule-binding domain to a low-affinity state [15, 17, 20, 51]. ATP hydrolysis at AAA3 and ATP binding at AAA4 regulate the AAA1-dependent release of dynein from microtubule [22, 52, 55-59]. Most relevant to our current study, structural studies combined with biophysical analyses in vitro on purified yeast dynein motor domain have provided significant insights into the function of the AAA3 ATPase cycle [22, 58, 59]. Specifically, when the AAA3 ATP hydrolysis is blocked, the linker fails to bend in response to ATP binding at AAA1 and dynein fails to release from microtubule [22, 58, 59]. It has been thought that the AAA3 ATPase cycle may participate in the spatial regulation of dynein in vivo [22, 58]; For example, the strong microtubule binding of dynein with its ATP-bound AAA3 may help anchor dynein at the vicinity of the microtubule plus end before the initiation of minus-end-directed movement. However, this interesting idea has not yet been examined in vivo in the context of cargo adapter, dynactin and another important dynein regulator LIS1. LIS1 deficiency causes human lissencephaly, and LIS1 is a dynein regulator in many organisms and cell types [6, 48, 60]. LIS1 binds to the dynein motor ring around AAA3/AAA4 and to a site on the stalk [61-64]. Different AAA3 nucleotide states cause the yeast dynein motor domain to respond to LIS1 differently in vitro: While LIS1 strongly inhibits the motility of apo-AAA3 dynein, it enhances the motility of ATP-AAA3 dynein [61]. However, as the relationship between LIS1 and the AAA3 ATPase cycle has not yet been examined in the context of dynactin and cargo adapters, it would be important to do so especially given the more recent knowledge of LIS1f ñ ’s involvement in dynein activation in the context of dynactin and cargo adapters [62, 65-70].

We have been studying the in vivo regulation of dynein using the filamentous fungus *Aspergillus nidulans* as a genetic system. Dynein in filamentous fungi is required for nuclear distribution [71-74] and for the transport of early endosomes and other cargoes, some of which move by hitchhiking on the motile early endosomes [75-85]. Inside the fungal hypha, dynein, dynactin and LIS1 all accumulate at the microtubule plus ends near the hyphal tip, and the plus-end accumulation of dynein and dynactin depends on kinesin-1 [76, 86-90]. The microtubule plus-end accumulation of dynein is important for dynein-mediated transport of early endosomes away from the hyphal tip [76, 91, 92]. The physical interaction between dynein and early endosome depends on the dynactin complex as well as the FHF (Fts, Hook and Fhip) complex (note that the FHF complex was first identified in mammalian cells [93]) [33-36, 94]. Within the *A. nidulans* FHF complex, HookA lacking the C-terminal cargo-binding domain (ΔC-HookA) interacts with dynein-dynactin [36]; FhipA mediates the interaction between HookA’s C-terminus and early endosome [94], most likely via its direct interaction with Rab5 on early endosome [95]. Recently, we found that ΔC-HookA overexpression drives the relocation of dynein and dynactin from the microtubule plus ends to the minus ends at septa or the spindle-pole bodies (SPBs) [68, 96], both of which contain active microtubule-organizing centers [97, 98]. This relocation is consistent with cargo-adapter-mediated dynein activation observed in vitro [24, 42, 65, 67]. By using this ΔC-HookA-driven dynein relocation as a readout for dynein activation in vivo, we found that dynein activation requires NudF/LIS1 in *A. nidulans* [68]. Importantly, by making the phi mutation preventing the formation of the phi structure [32], the requirement for NudF/LIS1 is partially bypassed, suggesting that LIS1 promotes the open state of dynein [68]. This conclusion is largely consistent with other recent studies [62, 69, 70, 99].

*A. nidulans* genetics has previously revealed a NudF/LIS1-bypass suppressor mutation in the AAA4 arginine finger of the dynein heavy chain [63, 100, 101], which is implicated in AAA3 ATP hydrolysis based on the knowledge from other AAA ATPases [102, 103]. Here we show that the AAA4 arginine finger mutation allows the function of NudF/LIS1 in the ΔC-HookA-driven dynein activation to be bypassed. We further created dynein AAA3 ATP-binding (Walker A or wA) and ATP-hydrolyzing (Walker B or wB) mutants and analyzed them in the presence and absence of dynactin, NudF/LIS1 or HookA. Surprisingly, the wB-AAA3 dynein accumulates strongly at the microtubule minus ends at septa even without the overexpression of ΔC-HookA. The wA-AAA3 dynein mainly accumulates at the hyphal tip but also shows a septal accumulation. The minus-end localization of wB-AAA3 dynein depends on the functionality of dynein as well as the dynactin complex, but it does not need NudF/LIS1 or HookA, suggesting that blocking AAA3 ATP hydrolysis would allow the requirement of LIS1 or HookA for dynein activation to be bypassed. The wA-AAA3 mutation has a relatively milder effect on bypassing the requirement of LIS1 or HookA, but it alters the dynamic nature of the dynein-LIS1 interaction, which will be discussed in more detail. Together, our data suggest that AAA3 ATPase cycle is critical for dynein regulation in vivo, and most importantly, AAA3 ATP binding modulates the LIS1-dynein interaction, and AAA3 ATP hydrolysis is needed for preventing dynein from being abnormally activated before binding its early endosome cargo.

## Results

### Dynein with the AAA4 arginine-finger mutation bypasses the requirement of NudF/LIS1 for ΔC-HookA-driven activation

In *A. nidulans*, genetic studies led to the identification of a *nudF (lis1)* bypass suppressor in dynein HC, *nudA*^R3086C^, which represents a mutation in the AAA4 arginine finger [63, 100, 101]. Here we tested if the AAA4 arginine finger (RF-AAA4) mutation allows the requirement of NudF/LIS1 for dynein activation to be bypassed. In the strain carrying the RF-AAA4 mutation, GFP-tagged dynein heavy chain (called GFP-dynein for simplicity) localizes along microtubules with a clear plus-end enrichment (Figure 1A) [100]. In addition, septal (minus-end) accumulation of dynein was also readily noticed in some hyphae (Figure 1A). Upon overexpression of ΔC-HookA (*gpdA*-ΔC-*hookA*-S), the RF-AAA4 dynein accumulates strongly at the septal minus ends (Figure 1A), similar to that exhibited by the wild-type dynein [68], suggesting that dynein with the RF-AAA4 mutation is activated by the cargo adapter. We then introduced the temperature-sensitive loss-of-function mutation of *nudF*/*lis*1, *nudF*6, to test if NudF/LIS1 is needed for this activation. Interestingly, a strong septal accumulation of the RF-AAA4 dynein was observed in the *nudF*6, *gpdA*-ΔC-*hookA*-S background (Figure 1A), which is in sharp contrast to the plus-end accumulation of wild-type dynein in the *nudF*6, *gpdA*-ΔC-*hookA*-S background (Figure 1A, B) [68]. Thus, the RF-AAA4 mutation allows the requirement of NudF/LIS1 for dynein activation to be bypassed.

**Figure 1.**
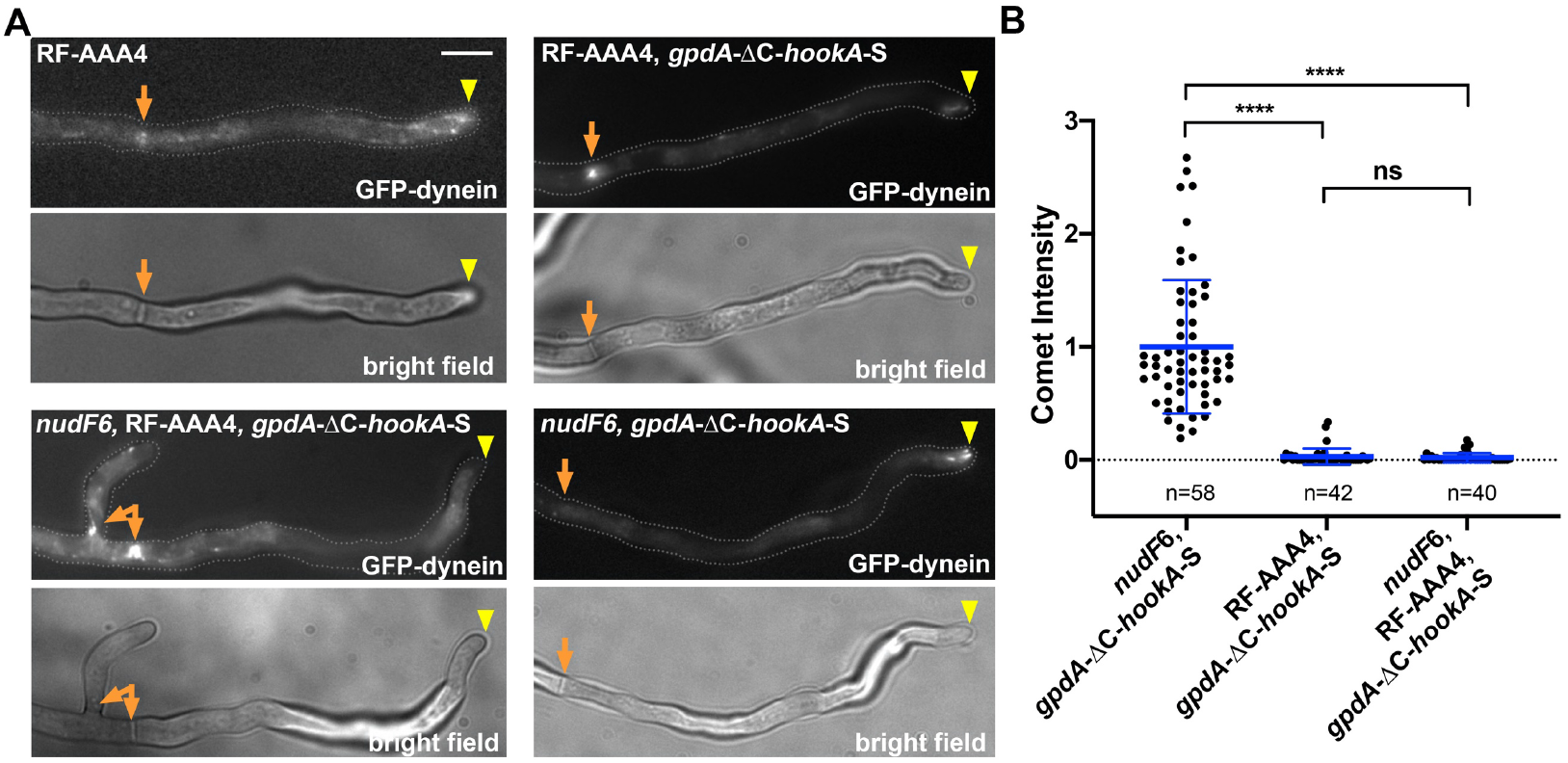
The AAA4 arginine-finer mutation (RF-AAA4 or *nudA*^R3086C^) allows NudF/LIS1 function to be bypassed during ΔC-HookA-mediated dynein activation. (A) Images of GPF-dynein are shown for a strain containing RF-AAA4, a strain containing RF-AAA4, *gpdA*-ΔC-*hookA*-S, a strain containing *nudF*6, RF-AAA4, *gpdA*-ΔC-*hookA*-S, and a strain containing *nudF*6, *gpdA*-ΔC-*hookA*-S. The strain containing *nudF*6, *gpdA*-ΔC-*hookA*-S was used as a control, as it shows plus-end comets formed by wild-type dynein. Bright-field images are shown below to indicate hyphal shape and position of septum. Hyphal tip is indicated by a yellow arrowhead and septum by a light brown arrow. Bar, 5 μm. (B) A quantitative analysis on dynein comet intensity in the *nudF*6, *gpdA-*ΔC-*hookA*-S strain (n=58), the RF-AAA4, *gpdA-*ΔC-*hookA*-S strain (n=42) and the *nudF*6, RF-AAA4, *gpdA-*ΔC-*hookA*-S strain (n=40). The average value for the *nudF*6, *gpdA-*ΔC-*hookA*-S strain is set as 1. Scatter plots with mean and S.D. values were generated by Prism 8. ****p<0.0001; ns, non-significant or p>0.05 (Kruskal-Wallis ANOVA test with Dunn’s multiple comparisons test, unpaired).

### Dynein function in vivo is affected by AAA3 ATP-binding or hydrolysis mutation; the AAA3 ATP-hydrolysis mutation suppresses the Δ*nudF/lis1* colony phenotype

The AAA4 arginine finger is implicated in ATP hydrolysis at AAA3 based on studies of other AAA+ ATPases [103]. Indeed, mutations in both the AAA4 arginine finger and the AAA3 Walker B motif affecting ATP-hydrolysis cause a significant reduction in dynein velocity in vitro [55, 57, 63]. Thus, we sought to directly analyze the role of the AAA3 ATPase cycle in dynein activation in vivo mediated by dynactin, LIS1 and cargo adapter. Based on the similarity of dynein heavy chain sequences from mammalian dynein and *A. nidulans* dynein (Figure S1), we constructed the AAA3 ATP-binding (Walker A or wA-AAA3, *nudA*^K2599A^) and hydrolyzing (Walker B or wB-AAA3, *nudA*^E2663Q^) mutants. Both of these mutants formed small colony with a defect in asexual spore production (indicated by the lack of colony color) (Figure 2A), consistent with a defect in nuclear migration to the spores [104]. In comparison to the RF-AAA4 mutant, which formed a colony with asexual spores (note that the green color of the colony is from the asexual spores), these AAA3 mutants are much sicker (Figure 2A). Thus, although the AAA4 arginine finger is implicated in AAA3 ATP hydrolysis, our data showed that it is not essential for this process in *A. nidulans*. Nevertheless, the colonies formed by the wA-AAA3 and wB-AAA3 mutants are obviously healthier than the Δ*nudA* (dynein heavy chain) [105] and the Δ*nudF* (*lis1*) [101] mutants colonies (Figure 2A). Thus, the wA-AAA3 or wB-AAA3 mutant dynein is partially functional in vivo. Consistently, both the wA-AAA3 and wB-AAA3 mutants exhibited a partial defect in nuclear distribution (Figure 2B, 2C). Importantly, the wB-AAA3 mutation clearly suppressed the colony growth defect of the Δ*nudF* mutant, suggesting that the wB-AAA3 mutation allows the function of NudF/LIS1 to be bypassed (Figure 2D). The wA-AAA3 mutation also made the Δ*nudF* colony slightly bigger at 32°C but the suppression is not as strong as that caused by the wB-AAA3 mutation (Figure 2D). Surprisingly, adding the *nudF*6 temperature-sensitive mutation to the wA-AAA3 mutant makes the wA-AAA3 mutant grew better at *nudF*6’s semi-permissive temperature of 32°C (Figure 2D). Because the *nudF*6 mutation makes NudF/LIS1 protein less stable [104], this result suggests that a decrease in the NudF/LIS1 protein level alleviates the defect caused by the wA-AAA3 mutation.

**Figure 2.**
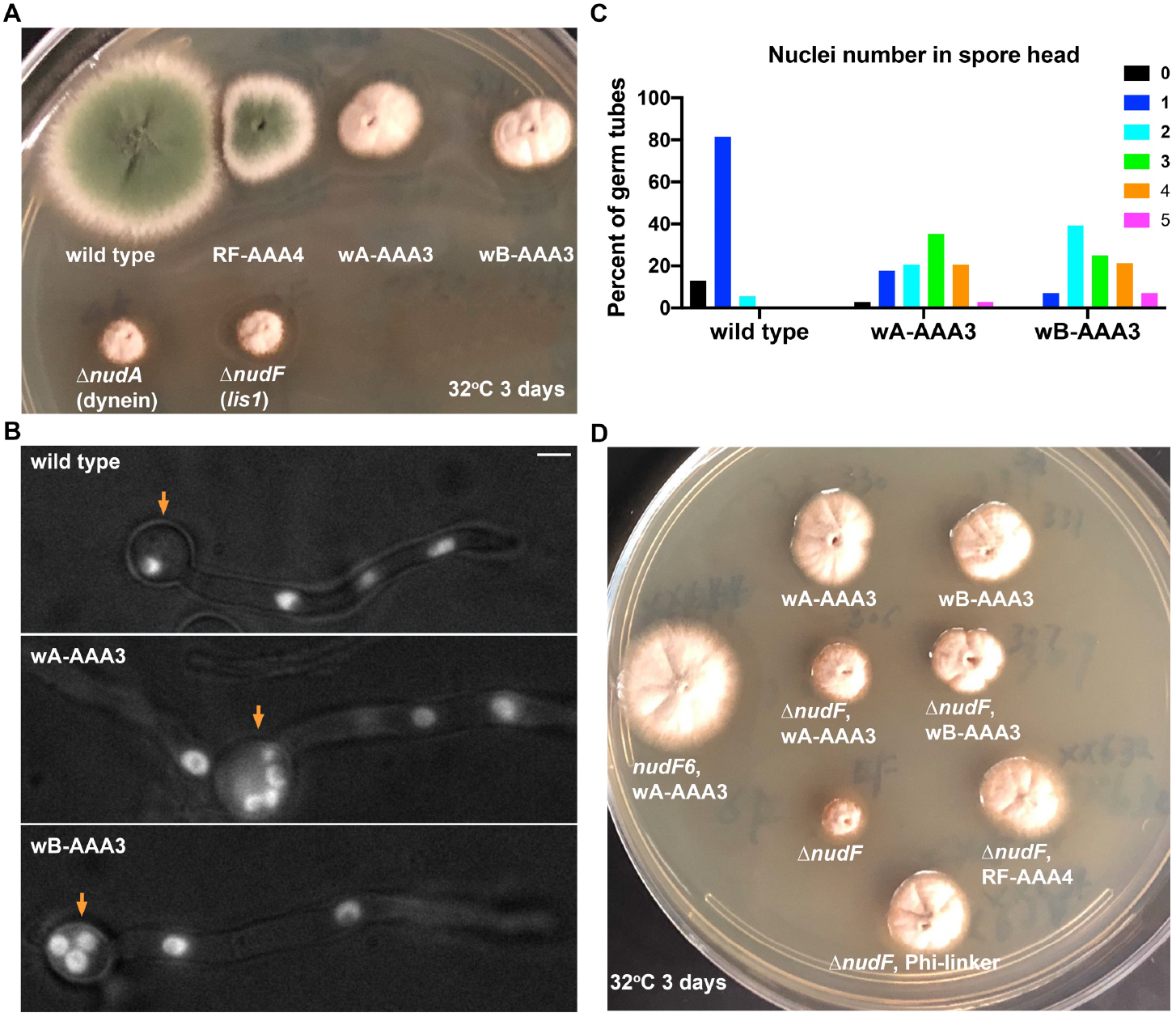
Both the wA-AAA3 and wB-AAA3 mutations affect dynein function, and the wB-AAA3 mutation suppresses the Δ*nudF* (lis1) growth defect. (A) Colony phenotypes of the wA-AAA3 and wB-AAA3 mutants in comparison with the wild type, RF-AAA4, Δ*nudA* and Δ*nudF* strains. The plate was incubated at 32°C for 3 days. (B) Images of nuclei labeled with NLS-DsRed in wild type, the wA-AAA3 mutant and the wB-AAA3 mutant. Spore head is indicated by a brown arrow. Bar, 5 μm. (C) A quantitative analysis on the percentage of germ tubes containing 0, 1, 2, 3, 4 or 5 nuclei in the spore head (wild type: n=54; wA-AAA3: n=34; wB-AAA3: n=28). The number of nuclei in the spore head of the wA-AAA3 mutant or the wB-AAA3 mutant is higher than that in wild type (p<0.0001 in both cases, Kruskal-Wallis ANOVA test with Dunn’s multiple comparisons test, unpaired). However, the number of nuclei in the spore head of the wA-AAA3 mutant is not significantly different from that in the wB-AAA3 mutant (p>0.99, Kruskal-Wallis ANOVA test with Dunn’s multiple comparisons test, unpaired). (D) Colony phenotypes of various single and double mutant strains. The plate was incubated at 32°C for 3 days.

### The minus-end localization of dynein at septa is enhanced mildly by the wA-AAA3 mutation and more dramatically by the wB-AAA3 mutation

We next used live cell imaging to observe the localization of GFP-dynein [100], carrying either the wA-AAA3 or wB-AAA3 mutation. We also observed early endosomes in the same hyphae labeled by mCherry-RabA [89, 91, 106]. In wild-type hyphae, dynein mainly accumulates at the microtubule plus ends near the hyphal tip as comet-like structures [86, 107], although localization of dynein at the septa (minus ends) is also seen [98, 108]. The wA-AAA3 mutant dynein signals are seen near the hyphal tip, although in some hyphae the signals do not look like typical plus-end comets and seem partially overlap with early endosomes abnormally accumulated at the hyphal tip (Figure 3A). In addition, the wA-AAA3 dynein signals also form a faint microtubule decoration with an enrichment at septa (Figure 3A). In comparison, the wB-AAA3 mutant dynein shows no hyphal tip-enriched signals but a more prominent accumulation at septa and microtubule decoration (Figure 3A, 3B). In some hyphae, the wB-AAA3 dynein is also accumulated as bright dots, which correspond to the SPB signals near the individual nuclei (Figure S2). In case of the wild-type dynein, such a strong septal and SPB accumulation was only observed when ΔC-HookA was overexpressed to drive dynein activation [68]. As the SPB signals are cell cycle-specific [96], we focus only on the septal accumulation that is more consistently observed. Previously, we showed that the phi mutant dynein (open dynein) also accumulates at the septa without the overexpression of ΔC-HookA, and it is co-localized with early endosomes at the septa [68]. However, early endosomes in either the wB-AAA3 mutant or the wA-AAA3 mutant exhibited a hyphal-tip accumulation instead of a septal accumulation (Figure 3A), although in some wB-AAA3 hyphae, we could still observe faint septal signals of early endosomes (Figure 3A). These observations suggest that most mutant dynein molecules go to the microtubule minus ends without its early endosome cargo.

**Figure 3.**
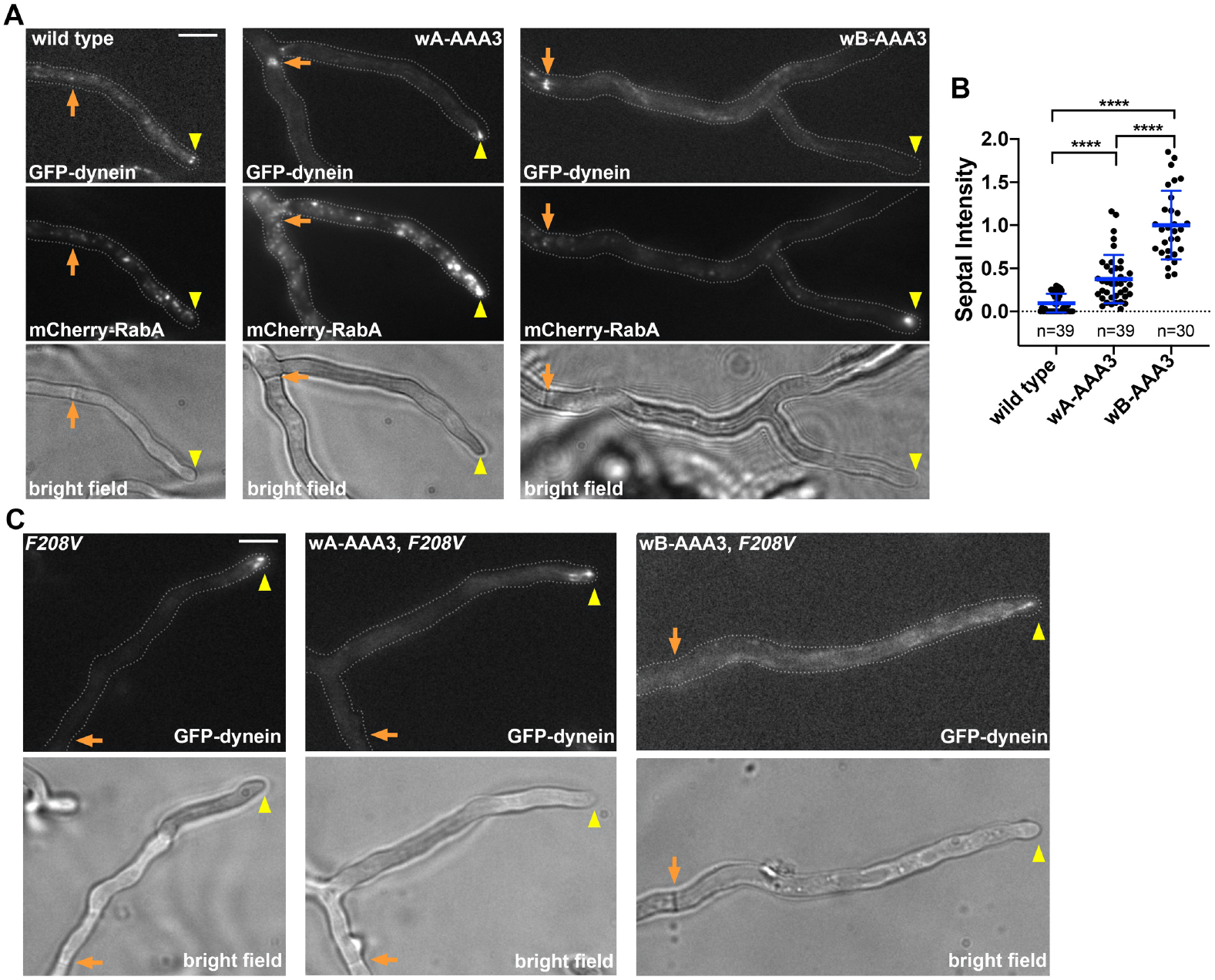
The wA-AAA3 or wB-AAA3 dynein partially accumulates at the microtubule minus ends at septa, and this accumulation requires dynein functionality. (A) Localization of GFP-dynein and early endosomes (mCherry-RabA) in wild type, the wA-AAA3 mutant and the wB-AAA3 mutant. Bright-field images are shown below to indicate hyphal shape and position of septum. Hyphal tip is indicated by a yellow arrowhead and septum by a light brown arrow. (B) A quantitative analysis on septal dynein intensity in wild type (n=39), wA-AAA3 (n=39), and wB-AAA3 (n=30) strains. The average value for the wB-AAA3 strain is set as 1. Scatter plots with mean and S.D. values were generated by Prism 8. ****p<0.0001, ***p<0.001, **p<0.01 (Kruskal-Wallis ANOVA test with Dunn’s multiple comparisons test, unpaired). (C) The dynein tail mutation *nudA*F^208V^ diminishes the septal accumulation of the wA-AAA3 or wB-AAA3 mutant dynein. Bright-field images are shown below to indicate hyphal shape and position of septum. Hyphal tip is indicated by a yellow arrowhead and septum by a light brown arrow. Bars, 5 μm.

Next, we examined the localization of dynactin in the wA-AAA3 and wB-AAA3 mutants by introducing the p150-GFP fusion into strains containing the mutations. In the wA-AAA3 mutant, the pattern of p150-GFP localization is similar to that of dynein localization, with both hyphal-tip and septal signals (Figure S3). In the wB-AAA3 mutant, the localization pattern of p150-GFP is also similar to that of wB-AAA3 dynein with a strong septal accumulation, although the microtubule decoration appears weaker, and faint plus-end comets were also observed in some hyphal tips. These observations suggest that dynactin is associated with dynein in these mutants, although some individual dynein and dynactin complexes are also present.

Although the wB-AAA3 dynein decorates microtubules besides forming the minus-end accumulation, the microtubule-bound dynein signals are mainly non-motile. Thus, the minus-end accumulation must have been established during many hours of hyphal growth. To address whether the full-length wB-AAA3 dynein is motile, we prepared cell extract from a strain containing GFP-labeled wB-AAA3 dynein for an in vitro motility assay. A strain containing GFP-labeled wild-type dynein was used as a positive control. In this control strain, the *gpdA*-ΔC-*hookA*-S allele was present to ensure that dynein mainly moves towards the microtubule minus end. We found that wild-type dynein moved robustly in the motility assay, and the full-length wB-AAA3 dynein also moves processively along the microtubule (when 3mM ATP instead of 1mM ATP was used), albeit with a very low speed (Figure S4A, S4B and S4C), consistent with previous in vitro data obtained from the yeast dynein motor domain [22, 52, 57-59, 61]. This very slow motility was also observed in vivo for the septum-directed movement of p150-GFP, which was detected very occasionally (Figure S4D), and it is also consistent with the notion that the wB-AAA3 dynein is partially functional in vivo (Figure 2).

In addition, we performed in vivo experiments to show that microtubules are important for the septal accumulation of the wB-AAA3 dynein. First, we treated the wB-AAA3 mutant with benomyl, a microtubule-depolymerizing drug. This treatment caused a significant decrease in the septal accumulation and a significant increase in the hyphal-tip accumulation of the wB-AAA3 dynein (Figure S5A-D). The signals of hyphal-tip dynein partially overlap with those of abnormally accumulated early endosome (Figure S5B), suggesting that the cytosolic wB-AAA3 dynein molecules can associate with early endosomes. We then replaced the benomyl-containing medium with regular medium to let the cells recover. The recovery caused a significance increase in the septal accumulation but a significant decrease in the hyphal-tip accumulation of the wB-AAA3 dynein compared to cells treated with benomyl. During the recovery, we were able to capture some transient dynein movements away from the hyphal tip (plus ends) (Figure S5E), further supporting the idea that the wB-AAA3 dynein can move towards the minus ends in live cells.

Next, we sought to use a molecular genetic approach to confirm that the septal (minus-end) accumulation of the AAA3 mutant dynein needs dynein functionality. To do that, we constructed GFP-tagged dynein heavy chain containing an AAA3 mutation and the *nudA*^F208V^ dynein tail mutation that significantly impairs dynein activity but does not impair the dynein-dynactin interaction or dynein-early endosome interaction [68, 109]. Dynein carrying the wA-AAA3 and *nudA*^F208V^ double mutations formed plus-end comets near hyphal tip just like dynein with the *nudA*^F208V^ single mutation (Figure 3C). Dynein carrying the wB-AAA3 and *nudA*^F208V^ mutations showed a microtubule decoration with a clear plus-end enrichment (Figure 3C). Importantly, the *nudA*^F208V^ mutation diminished the minus-end accumulation of the mutant dynein, as the septal accumulation disappeared in both the wA-AAA3, *nudA*^F208V^ and the wB-AAA3, *nudA*^F208V^ double mutants (Figure 3C). This result confirmed that the minus-end accumulation in both the wA-AAA3 or wB-AAA3 single mutants requires dynein functionality.

### Activation of wA-AAA3 dynein depends largely on HookA, but activation of wB-AAA3 dynein occurs without HookA, although it still needs dynactin

In *A. nidulans*, the early endosomal adapter HookA is a critical dynein activator, and overexpression of the cytosolic ΔC-HookA drives dynein relocation from the microtubule plus ends to the minus ends [68]. We suspected that activation of the AAA3 mutant dynein could be HookA-independent because the HookA-bound early endosomes did not co-localize with dynein at the septa. However, it would be hard to exclude the possibility that the mutant dynein is activated by the HookA-bound early endosome and then dissociates abnormally from the early endosome. To directly test whether HookA is needed for activating the wA-AAA3 and wB-AAA3 dynein, we introduced the *gpdA*-ΔC-*hookA*-S allele (causing overexpression of ΔC-HookA) [68] or the Δ*hookA* allele [36] into strain background with GFP-dynein harboring the wA-AAA3 or wB-AAA3 mutation. We found that activation of the wA-AAA3 dynein is largely but not absolutely HookA-dependent. Overexpression of ΔC-HookA (*gpdA*-ΔC-*hookA*-S) drives the septal accumulation of the wA-AAA3 dynein, just like wild-type dynein (Figure 4A). In the Δ*hookA* mutant, wild-type dynein forms strong comets near the hyphal tip, and the wA-AAA3 dynein forms plus-end comets as well but also a microtubule decoration and a septal enrichment (Figure 4B). As a consequence, the wA-AAA3 dynein’s plus-end signal intensity is lower than that of wild-type dynein in the Δ*hookA* background (Figure 4C). These results suggest that wA-AAA3 dynein needs HookA for activation just like wild-type dynein, but a small portion of the wA-AAA3 dynein molecules may be abnormally activated in a HookA-independent fashion. On the other hand, the wB-AAA3 dynein shows a strong septal accumulation not only upon overexpression of ΔC-HookA (*gpdA*-ΔC-*hookA*-S) but also in the Δ*hookA* background (Figure 4A, 4B). The septal signal intensity of the wB-AAA3 dynein in the Δ*hookA* background is significantly higher than that of the wild-type dynein or the wA-AAA3 dynein in the Δ*hookA* background (Figure 4D). Moreover, while loss of HookA clearly decreases the septal signal intensity of the wA-AAA3 dynein (Figure 4E), it does not significantly decrease the septal signal intensity of the wB-AAA3 dynein (Figure 4F). Thus, the abnormal HookA-independent dynein activation occurs at a much higher degree in the wB-AAA3 mutant than in the wA-AAA3 mutant. However, while the microtubule decoration by the wB-AAA3 dynein was obvious in the Δ*hookA* background, overexpression of ΔC-HookA in the *gpdA*-ΔC-*hookA*-S background drives almost all the wB-AAA3 mutant dynein to septa, indicating that HookA still can facilitate the activation of wB-AAA3, despite the tendency of the mutant dynein to undergo HookA-independent activation.

**Figure 4.**
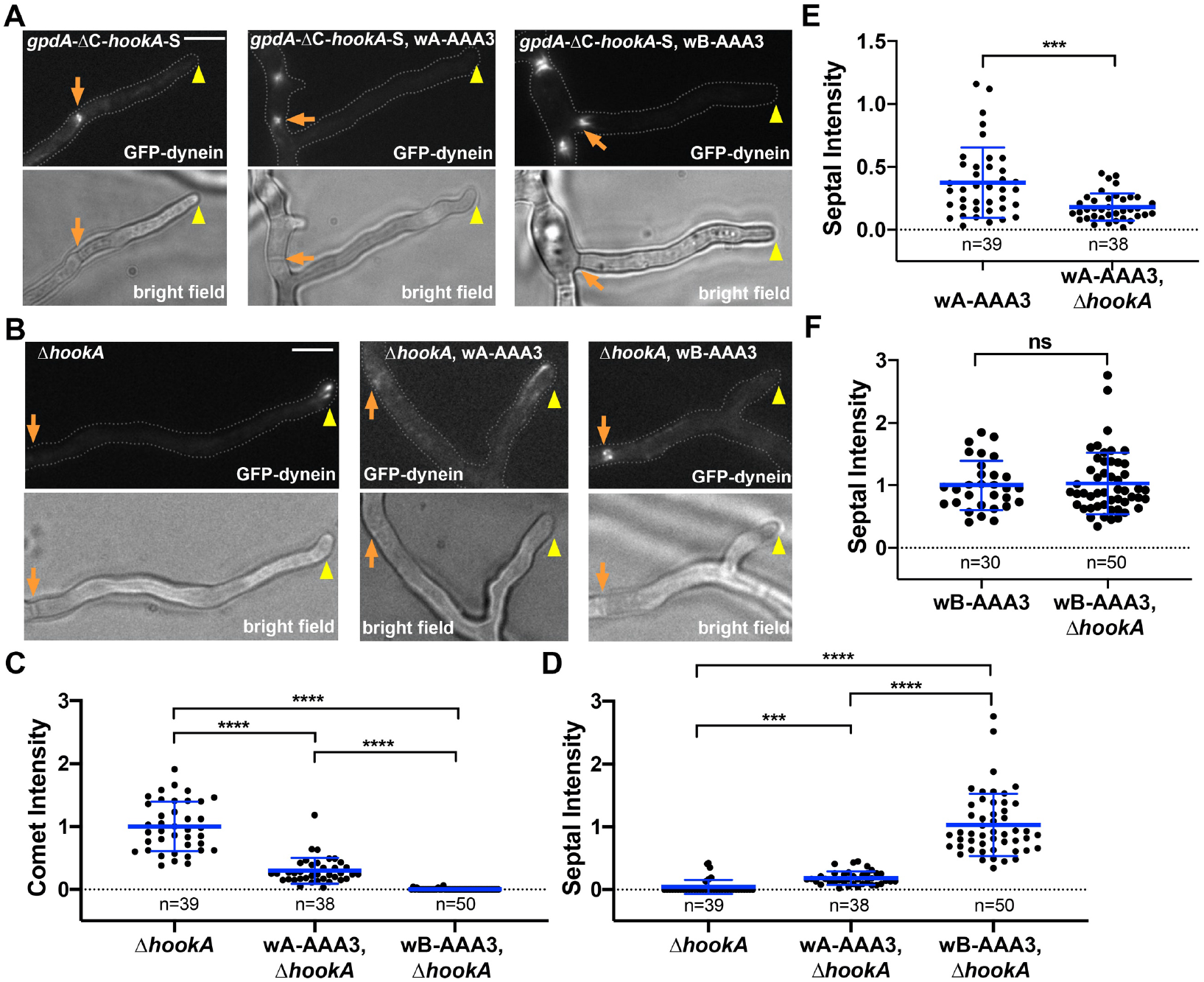
Different effects of the cargo adapter HookA on the localization of wild-type dynein, wA-AAA3 dynein and wB-AAA3 dynein. (A) Wild-type dynein, wA-AAA3 dynein and wB-AAA3 dynein in the *gpdA*-ΔC-*hookA*-S (overexpression of ΔC-HookA) background. Bright-field images are shown below to indicate hyphal shape and position of septum. Hyphal tip is indicated by a yellow arrowhead and septum by a light brown arrow. Bar, 5 μm. (B) Wild-type dynein, wA-AAA3 dynein and wB-AAA3 dynein in the Δ*hookA* mutant background. Bright-field images are shown below to indicate hyphal shape and position of septum. Hyphal tip is indicated by a yellow arrowhead and septum by a light brown arrow. (C)(D) Quantitative analyses on comet intensity and septal intensity of wild type (n=39), wA-AAA3 (n=38), and wB-AAA3 (n=50) dynein in the Δ*hookA* background. Scatter plots with mean and S.D. values were generated by Prism 8. ****p<0.0001, ***p<0.001 (Kruskal-Wallis ANOVA test with Dunn’s multiple comparisons test, unpaired). In (C), the average value for wild-type dynein in the Δ*hookA* strain is set as 1. In (D), the average value for wB-AAA3, Δ*hookA* strain is set as 1. (E) A quantitative analysis on septal intensity of wA-AAA3 dynein in the wild-type (n=39) and the Δ*hookA* (n=38) backgrounds. (F) A quantitative analysis on septal intensity of wB-AAA3 in the wild-type (n=30) and the Δ*hookA* (n=50) backgrounds. In (E) and (F), the average value for the wB-AAA3 dynein in the wild-type background is set as 1. Scatter plots with mean and SD values were generated by Prism 8. ***p<0.001; ns, non-significant or p>0.05 (unpaired, Mann-Whitney test, Prism 8).

The dynactin complex is a critical component in dynein activation, as the Arp1 mini-filament provides the dynein tail binding sites [24-28, 32]. However, this function of dynactin has not been directly addressed in *A. nidulans* due to its additional role in the microtubule plus-end accumulation of wild-type dynein. To determine whether dynactin is required for activating the wA-AAA3 or wB-AAA3 dynein, we introduced the Arp1 conditional-null (*alcA*-*nudK*^Arp1^) allele [110] into the strain background with GFP-dynein harboring the wA-AAA3 or wB-AAA3 mutation. In the background of wild-type dynein, shutting down Arp1 expression using this *alcA*-*nudK*^Arp1^ allele causes a loss of microtubule plus-end accumulation of dynein [110]. Since the *alcA* promoter is repressed by glucose but de-repressed by glycerol [111], we used both glycerol- and glucose-containing media to culture the cells containing wA-AAA3, *alcA*-*nudK*^Arp1^ and wB-AAA3, *alcA*-*nudK*^Arp1^ double mutants. In the single AAA3 mutants without the *alcA*-*nudK*^Arp1^ allele, the same localization patterns were observed under these two culture conditions (Figure 5A). In glycerol medium that allows the expression of Arp1 from the *alcA* promoter, the double mutants containing the *alcA*-*nudK*^Arp1^ allele showed the localization patterns similar to those in the single mutants. However, the double mutants showed drastically different localization patterns when Arp1 expression is shut off by glucose. Specifically, loss of Arp1 causes the wA-AAA3 dynein to become diffused in the cytoplasm, consistent with the previously identified role of dynactin in dynein’s plus-end accumulation [87, 88, 90, 107, 110]. For the wB-AAA3 dynein, loss of Arp1 allows it to more obviously decorate microtubules but significantly diminishes the septal (minus-end) accumulation (Figure 5A, B), indicating that dynactin is needed for activating the wB-AAA3 dynein.

**Figure 5.**
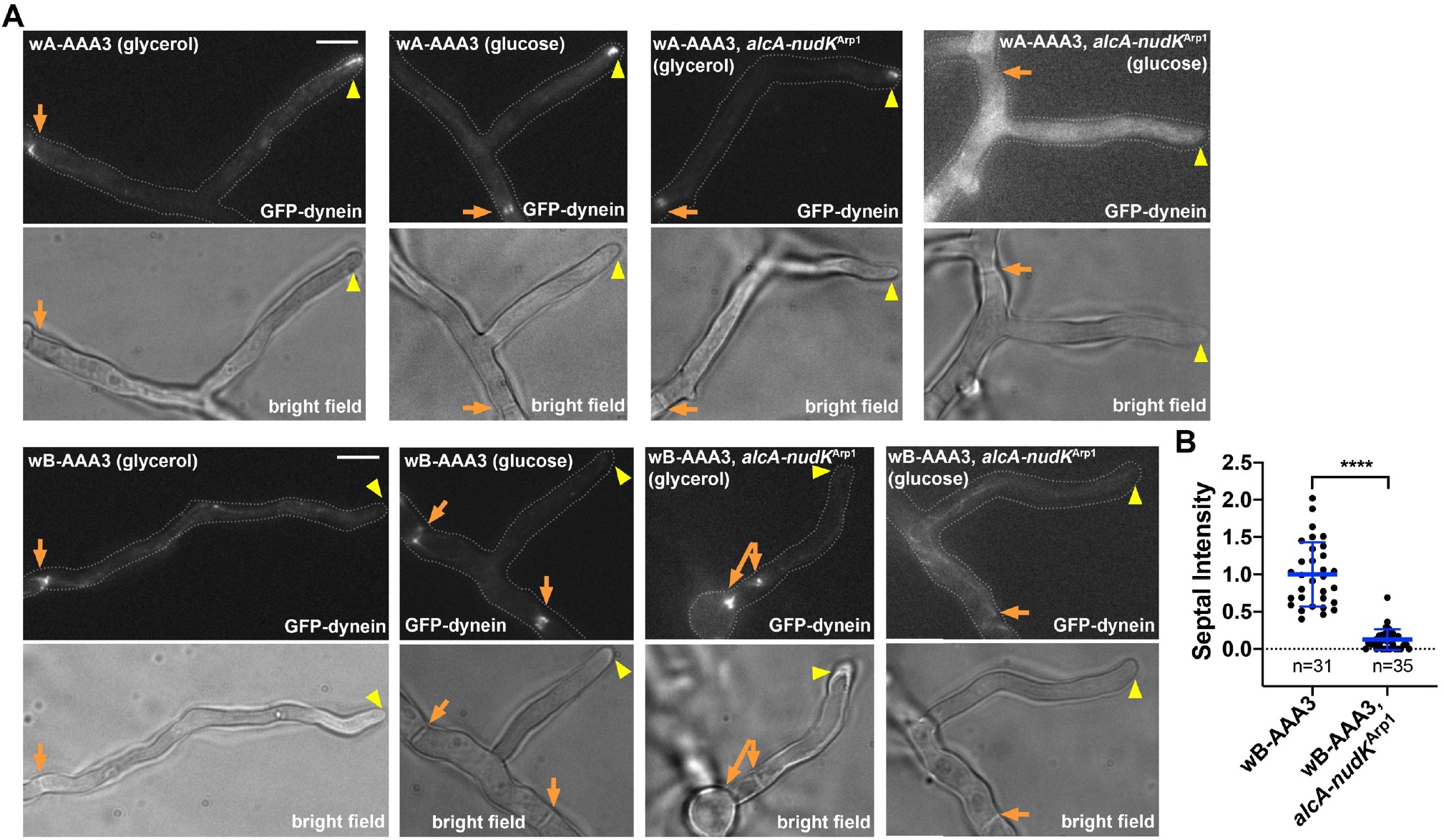
Dynactin is required for the septal accumulation of the wA-AAA3 or wB-AAA3 dynein. (A) Images of GFP-labeled wA-AAA3 dynein and wB-AAA3 dynein in the wild type or the Arp1 conditional-null (*alcA*-*nudK*^Arp1^) background. The *alcA* promoter is active on glycerol but repressed by glucose, and thus, the expression of Arp1 is shut off in strains containing the *alcA*-*nudK*^Arp1^ allele grown on glucose. The strains without the *alcA*-*nudK*^Arp1^ allele grown on glycerol or glucose were shown as controls. Bright-field images are shown below to indicate hyphal shape and position of septum. Hyphal tip is indicated by a yellow arrowhead and septum by a light brown arrow. Bar, 5 μm. (B) A quantitative analysis on septal intensity of the wB-AAA3 strain (n=31) and the wB-AAA3, *alcA*-*nudK*^Arp1^ strain (n=35) grown on glucose. Scatter plots with mean and SD values were generated by Prism 8. ****p<0.0001 (unpaired, Mann-Whitney test, Prism 8).

### Activation of the wB-AAA3 dynein does not need NudF/LIS1, but NudF/LIS1 is useful for activating the wA-AAA3 dynein

Next, we observed GFP-dynein harboring the wA-AAA3 or wB-AAA3 mutation in the Δ*nudF* mutant background. The wA-AAA3 dynein is totally accumulated at the microtubule plus ends as represented by bright comets in the absence of NudF/LIS1, suggesting that even the residual HookA-independent septal accumulation of the wA-AAA3 dynein is still dependent on NudF/LIS1. In contrast, the wB-AAA3 dynein accumulated strongly at septa regardless of whether NudF/LIS1 is present or not (Figure 6A). This result indicates that the requirement of NudF/LIS1 can be bypassed by the wB-AAA3 mutation, consistent with the strong suppression of the Δ*nudF* colony defect by the wB-AAA3 mutation (Figure 2D).

**Figure 6.**
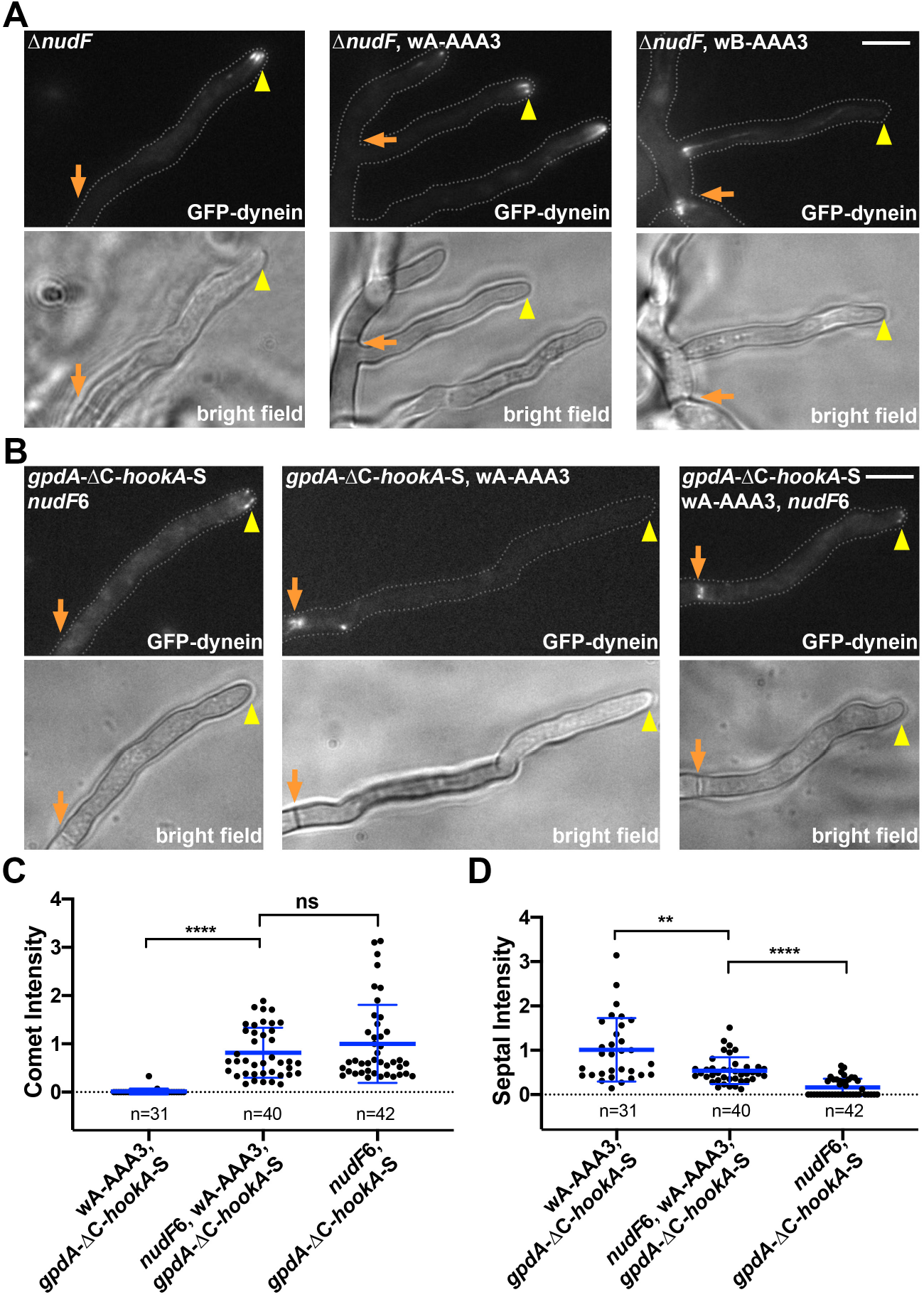
NudF/LIS1 is dispensable for the septal accumulation of the wB-AAA3 dynein but not the wA-AAA3 dynein, but overexpression of ΔC-HookA allows the wA-AAA3 mutation to partially bypass the requirement of NudF/LIS1. (A) Images of GFP-labeled wild-type dynein, wA-AAA3 dynein and wB-AAA3 dynein in the Δ*nudF* background. Bright-field images are shown below to indicate hyphal shape and position of septum. Hyphal tip is indicated by a yellow arrowhead and septum by a light brown arrow. Bar, 5 μm. (B) Images of GFP-labeled wA-AAA3 dynein or wild-type dynein in the *gpdA*-ΔC-*hookA*-S or the *nudF*6, *gpdA*-ΔC-*hookA*-S background. (C) (D) Quantitative analysis on comet intensity (C) and septal intensity (D) of the wA-AAA3 dynein in the *gpdA*-ΔC-*hookA*-S (n=31) or the *nudF*6, *gpdA*-ΔC-*hookA*-S background (n=40) and that of wild-type dynein in the *nudF*6, *gpdA*-ΔC-*hookA*-S background (n=42). Scatter plots with mean and SD values were generated by Prism 8. ****p<0.0001, **p<0.01; ns, non-significant or p>0.05 (unpaired, Mann-Whitney test, Prism 8).

Since the septal localization of wA-AAA3 dynein depends on NudF/LIS1, we further examined if overexpression of ΔC-HookA would help overcome this dependency. In this experiment, we used the temperature-sensitive *nudF*6 allele instead of the Δ*nudF* deletion allele to facilitate strain growth on solid medium before the live-cell imaging experiments. Previously, we found that while ΔC-HookA overexpression (*gpdA*-ΔC-*hookA*-S) causes dynein to relocate from the microtubule plus ends to the minus ends, the *nudF*6 mutation prevents this relocation and keeps dynein at the plus ends [68]. Consistent with a role of NudF/LIS1 in activation of the wA-AAA3 dynein, the *nudF*6 mutation also kept the wA-AAA3 dynein as plus-end comets upon overexpression of ΔC-HookA (*gpdA*-ΔC-*hookA*-S) (Figure 6B, 6C). However, a septal accumulation of the wA-AAA3 dynein was also obvious in the *nudF*6, *gpdA*-ΔC-*hookA*-S background, which was not observed for wild-type dynein (Figure 6B, 6D). Thus, the wA-AAA3 mutation bypasses NudF/LIS1 function to a low but detectable degree when ΔC-HookA is overexpressed. This may explain why the wA-AAA3 mutation mildly suppresses the colony growth defect of the Δ*nudF* mutant (Figure 2D).

### The wA-AAA3 mutation alters the LIS1-dynein interaction

Previously, we found that during the ΔC-HookA-mediated activation of wild-type dynein, NudF/LIS1 accumulates at the microtubule plus ends instead of going to the minus ends with dynein despite its importance for dynein activation [68]. This is consistent with LIS1’s dissociation from the activated dynein complex during minus-end-directed movement [62, 67, 69, 76, 88]. In the wB-AAA3 dynein mutant, although dynein strongly accumulates at the septal minus ends, NudF/LIS1-GFP forms plus-end comets instead of co-localizing with dynein at the septa, which is similar to the behavior of NudF/LIS1-GFP during the ΔC-HookA-driven activation of wild-type dynein (Figure 7A). Surprisingly, in the wA-AAA3 mutant, NudF/LIS1-GFP localizes to both the hyphal tip and septa just like wA-AAA3 dynein. Next, we introduced the *gpdA*-ΔC-*hookA*-S allele into the NudF/LIS1-GFP, wA-AAA3 strain and the NudF/LIS1-GFP, wB-AAA3 strain to examine NudF/LIS1-GFP localization upon overexpression of ΔC-HookA (Figure 7B). In the wA-AAA3, *gpdA*-ΔC-*hookA*-S background, where the wA-AAA3 dynein accumulates at the septa (Figure 4A), NudF/LIS1-GFP also accumulates at the septa (Figure 7B, 7C), although plus-end comets were still observed. In contrast, NudF/LIS1-GFP does not go to the septa in the wB-AAA3, *gpdA*-ΔC-*hookA*-S strain.

**Figure 7.**
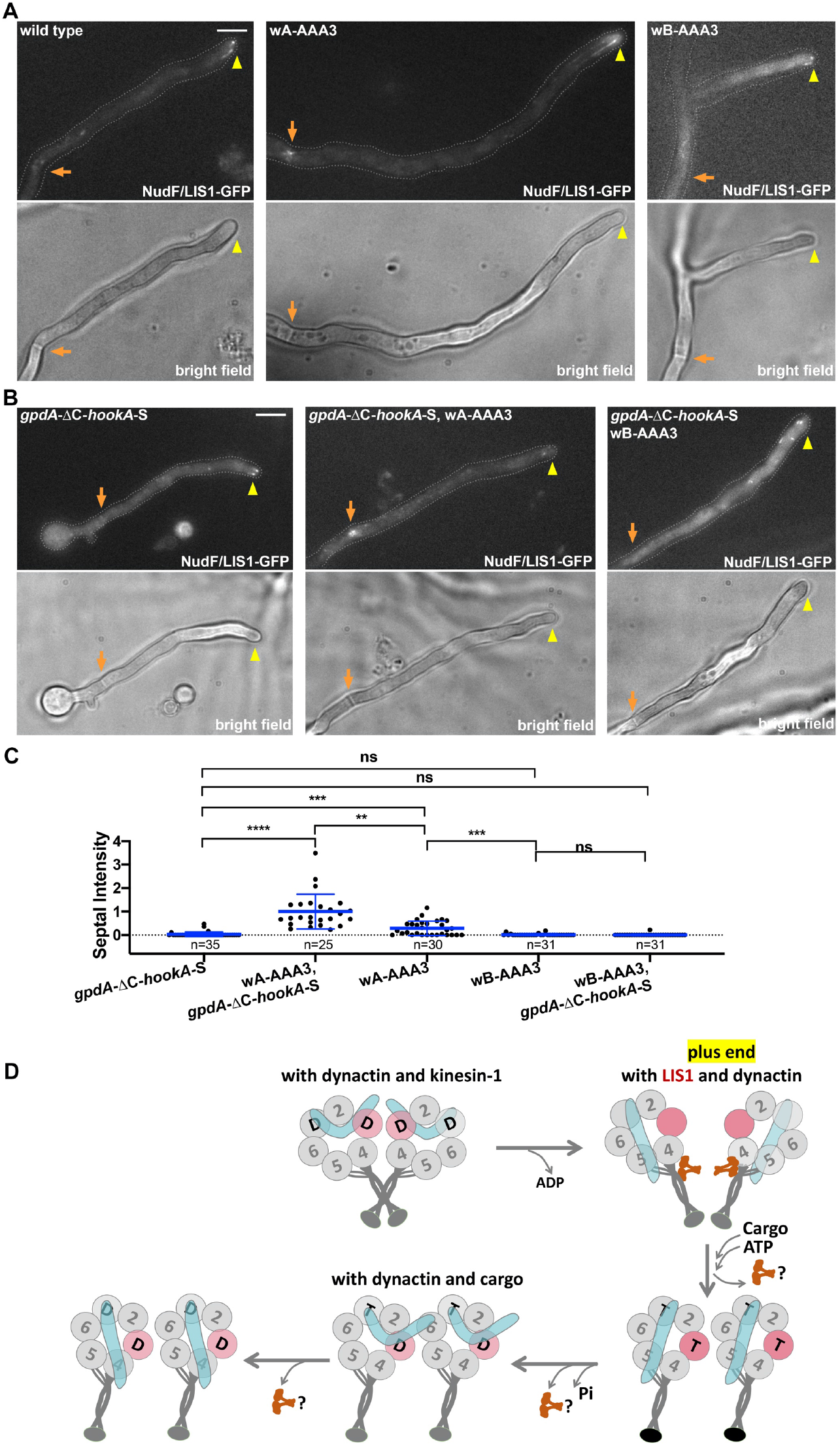
Images showing that NudF/LIS1 accumulates at the septa in the wA-AAA3 mutant and a hypothetic model of dynein activation involving the AAA3 ATPase cycle. (A) Images of NudF/LIS1-GFP localization in wild-type, the wA-AAA3 mutant and the wB-AAA3 mutant. Bright-field images are shown below to indicate hyphal shape and position of septum. Hyphal tip is indicated by a yellow arrowhead and septum by a light brown arrow. Bar, 5 μm. (B) Images of NudF/LIS1-GFP localization in wild-type, the wA-AAA3 mutant and the wB-AAA3 mutant in the *gpdA*-ΔC-*hookA*-S background. (C). A quantitative analysis on septal intensity of LIS1-GFP in the backgrounds of *gpdA*-ΔC-*hookA*-S (n=35), wA-AAA3, *gpdA*-ΔC-*hookA*-S (n=25), wA-AAA3 (n=30), wB-AAA3 (n=31), and wB-AAA3, *gpdA*-ΔC-*hookA*-S (n=31). The average value of the LIS1-GFP septal intensity in the wA-AAA3, *gpdA*-ΔC-*hookA*-S strain is set as 1. Scatter plots with mean and S.D. values were generated by Prism 8. ****p<0.0001, ***p<0.001, **p<0.01, ns, non-significant or p>0.05 (Kruskal-Wallis ANOVA test with Dunn’s multiple comparisons test, unpaired). (D) A hypothetic model of dynein activation involving the AAA3 ATPase cycle. Note that we only focus on a series of events at the vicinity of the microtubule plus end. During kinesin-1-mediated transport of dynein-dynactin to the microtubule plus end, dynein is most likely in the autoinhibited phi conformation. In the phi dynein, both AAA1 and AAA3 are occupied by ADP, and the linker is in a bent conformation (PDB code: 5NUG) [32]. Two different states (either apo or ATP at both AAA1 and AAA3) with the linkers docked on AAA5 [14, 22] are shown at the plus end. LIS1 may preferentially bind dynein with apo-AAA3, which is followed by cargo binding to cause dynein activation. ATP binding to AAA3 most likely happens after cargo-mediated dynein activation, because otherwise dynein would be activated by dynactin without the cargo. During the minus-end-directed movement of the dynein-dynactin-cargo adapter complex, AAA3 is ADP-bound most of the time, and the linker is flexible and its position would depend on the AAA1 nucleotide state [22, 58]. Finally, LIS1 can dissociate from the activated dynein-dynactin-cargo adapter complex at any step as long as the AAA3 is not in the apo state.

These results suggest that the wA-AAA3 mutation allows NudF/LIS1 to stay with dynein and co-localize with dynein at the microtubule minus ends, while the wB-AAA3 dynein tends to dissociate from NudF/LIS1 just like wild-type dynein after its cargo adapter-mediated activation.

## Discussion

Significant progress has been made in understanding dynein activation [2, 24, 32, 38, 42, 62, 68-70] and regulatory elements of the dynein motor domain [20, 112]. Most relevant to our current study is a detailed understanding on the role of AAA3 ATP hydrolysis in dynein’s mechanochemical cycle [22, 58, 59]. However, as the dynein motor mechanism is mainly studied in vitro by using the dynein motor domain itself, it has been unclear whether and how dynein’s mechanochemical cycle itself is linked to dynein activation in vivo. In this current study, we first found that an *A. nidulans* AAA4 arginine-finger mutation (implicated in AAA3 ATP hydrolysis) of dynein allows the function of NudF/LIS1 in dynein activation to be bypassed. This led us to further study the behaviors of wA-AAA3 (ATP-binding mutant) and wB-AAA3 (ATP-hydrolyzing mutant) dynein in the context of dynactin, the cargo adapter HookA and NudF/LIS1 in vivo. Our data suggest that the requirement for dynein activation is altered in both AAA3 mutants but more significantly in the wB-AAA3 mutant. In wild-type cells, dynein accumulates at the microtubule plus end and its activation needs both LIS1 and HookA, in addition to dynactin [49]. In contrast, the wB-AAA3 dynein can be activated by dynactin in the absence of LIS1 or HookA, as evidenced by its dynactin-dependent microtubule minus-end accumulation in the *nudF*/*lis*1 or *hookA* deletion mutant. Thus, AAA3 ATP hydrolysis is needed for preventing dynein from getting fully activated before binding its early endosome cargo. Moreover, our data also suggest that the wA-AAA3 mutation stabilizes the dynein-LIS1 interaction, suggesting that the dynamic nature of the dynein-LIS1 interaction, which has been observed both in vivo [76, 88, 113] and in vitro [62, 67, 69], is regulated by ATP binding at AAA3.

The wB-AAA3 dynein does not accumulate at the microtubule plus end, but rather, it shows a strong accumulation at the minus end, even in the absence of NudF/LIS1 or the cargo adapter HookA. The absence of the plus-end accumulation is most likely due to an unscheduled activation of the wB-AAA3 dynein, as its plus-end accumulation can be partially restored if dynein activity is compromised by a dynein tail mutation. The unscheduled activation may also weaken the kinesin-1-dynein interaction if kinesin-1 preferentially transports dynein in the autoinhibited phi conformation [49]; or, the mutant dynein may still bind kinesin-1 but overpowers it [114]. In the budding yeast, where dynein is exclusively required for spindle orientation [115], plus-end dynein accumulation depends on Pac1/LIS1, Bik1/Clip170 and Kip2/kinesin-7 [116-121]. While the full-length wA-AAA3 yeast dynein accumulates at the microtubule plus ends [120], the localization of the full-length wB-AAA3 dynein has yet to be studied in yeast. In filamentous fungi, LIS1 is not as important as dynactin for the plus-end accumulation of dynein [76, 87, 88, 90, 110]. This makes the activating function of LIS1 to be much more easily observed upon cargo-adapter-mediated dynein activation (compared to Num1-mediated activation in yeast [113]), as the lack of NudF/LIS1 prevents dynein from leaving the plus end [68]. In addition, while the *A. nidulans* AAA4 arginine-finger mutation or the wB-AAA3 mutation suppresses the growth defect of the Δ*nudF*/*lis1* mutant, the yeast AAA4 arginine-finger mutation does not suppress the Δ*pac*1/*lis1* mutant phenotype, because the additional function of Pac1/LIS1 in plus-end dynein accumulation cannot be bypassed [63]. In mammalian cells or if mammalian dynein is used in vitro, LIS1 also plays an important role in dynein’s plus-end accumulation (although dynactin is needed too) [47, 65, 67]. Thus, it may be easier to test if the mammalian wB-AAA3 dynein bypasses the requirement of LIS1 in an experimental setting where plus-end dynein accumulation is not a complicating factor. Despite the differences in the mechanisms of plus-end dynein localization, the dynein motor mechanism is most likely conserved in different organisms. Consistent with previous results on the yeast wB-AAA3 dynein motor domain [22, 52, 57-59, 61], the wB-AAA3 mutation also significantly inhibits the in vitro motility of full-length dynein in *A. nidulans* extract (Figure S4).

In this study, we found that the wB-AAA3 mutation allows dynein to bypass the requirement of NudF/LIS1, consistent with its suppression on the growth defect of the Δ*nudF* (lis1) mutant colony, an effect similarly caused by the AAA4 arginine-finger mutation or the phi mutation [63, 68, 100, 101]. As recent data link the function of LIS1 to promoting the open state of dynein [62, 68-70, 99], our current result suggest that the wB-AAA3 dynein is also open. In the wB-AAA3 mutant, the linker is locked at one position during the AAA1 ATPase cycle, with its N-terminus docked at AAA5 [22]. This is in contrast to wild-type dynein in which the linker bends upon ATP binding to AAA1, with its N-terminus close to AAA2/AAA3 and/or other positions [22]. The phi conformation of dynein is maintained by multiple interactions, and linkers in the phi dynein (PDB code: 5NUG) are bent [32]. Thus, if the linker is docked at AAA5, the phi structure may not be able to form, which implies that the wB-AAA3 dynein is unable to form the phi structure. However, the open conformation of the wB-AAA3 dynein must be different from that of the phi mutant dynein, which is open due to the weakened AAA4-linker ionic interaction between two dynein heavy chains within the dimer [32]. The phi mutant dynein also strongly accumulates at the microtubule minus ends at centrosomes in mammalian cells and at the septa in *A. nidulans* [32, 68]. However, a key difference between the phi mutant and the wB-AAA3 mutant is in that early endosomes strongly colocalize with the phi mutant (open) dynein [68] but not with the wB-AAA3 dynein at the septa. Although it is hard to completely exclude the possibility that other cargo adapters may be responsible for this minus-end localization, our data are most easily explained by the possibility that blocking AAA3 ATP hydrolysis allows the requirement of cargo adapter for dynein activation to be bypassed. The function of the cargo adapter has been implicated in enhancing the dynein-dynactin interaction [24, 42, 43, 47]. In fungi, however, the dynein-dynactin interaction clearly occurs before cargo-adapter-mediated dynein activation [48, 49], especially since dynactin is absolutely required for dynein’s plus-end accumulation in filamentous fungi [76, 87, 88, 90, 110] and dynein is required for the plus end-accumulation of dynactin in the budding yeast [122]. Thus, the function of the cargo adapter must be needed for changing the binding configuration of the two complexes so that the tails and motor domains of the dynein dimer can assume a parallel configuration [32]. Given the special linker position of the wB-AAA3 dynein [22], could the dynactin complex be sufficient for making the wB-AAA3 dynein parallel? In this context, we would also point out that in some previous in vitro experiments, dynactin alone can indeed cause or enhance dynein processivity in a moderate way [123-126]. Alternatively, could there be an in vivo regulator that reinforces the inactive state of dynein-dynactin before cargo binding and the wB-AAA3 mutation prevents dynein-dynactin from interacting with this regulator? Future work will clearly be needed to address these interesting questions.

LIS1 tends to dissociate from the dynein-dynactin-cargo adapter complex in fungal hyphae, budding yeast and in vitro assays [62, 67-69, 76, 88, 120], although it co-migrates with the dynein-dynactin-cargo adapter complex in other in vitro assays [65, 66]. In *A. nidulans*, the notion of LIS1-dynein dissociation after dynein activation explains the observation that LIS1 does not co-localize with activated dynein at the septal minus ends [68]. Interestingly, if AAA3 ATP binding is blocked, NudF/LIS1 strongly accumulate at the minus ends just like activated dynein, suggesting that the AAA3 ATPase cycle regulates the dynamic dynein-LIS1 interaction in vivo. It is possible that the abnormally increased dynein-LIS1 affinity contributes to the functional defect of the wA-AAA3 dynein, since reducing the NudF/LIS1 level in *A. nidulans* by using a *nudF*6 mutant grown at a semi-permissive temperature improves the growth of the wA-AAA3 mutant (Figure 2D). Based on this current study and previous studies from several labs, we propose that the AAA3 ATPase cycle is closely linked to the spatial regulation of dynein activation in vivo in the following ways: First, during kinesin-1-mediated transport to the microtubule plus end, dynein is in the autoinhibited phi conformation with an ADP-bound AAA3 (Figure 7D). Second, LIS1 may preferentially bind dynein with an apo-AAA3 (note that LIS1 binding is not compatible with the phi dynein conformation [62, 70]). Third, ATP binding to AAA3 occurs after cargo binding, as otherwise dynein will be activated by dynactin alone without the cargo. Finally, after cargo-mediated dynein activation, LIS1 may dissociate at any step except when AAA3 is in the apo state. We should point out that our current study is done in the context of not only cargo adapter and dynactin, but also NudE, another dynein regulator that binds both LIS1 and dynein [127-141]. Previous work on purified mammalian dynein showed that while LIS1 only binds dynein when ATP and vanadate are added, the inclusion of NudE allows LIS1-dynein binding in the absence of nucleotide [135]. Although the wA-AAA3 mutation clearly alters the dynamic LIS1-dynein interaction in *A. nidulans*, our hypothesis that LIS1 preferentially binds dynein with an apo-AAA3 before cargo binding will need to be further examined in the presence and absence of NudE.

In summary, our results from *A. nidulans* suggest that ATP binding and hydrolysis at AAA3 are important for different aspects of cargo-mediated dynein activation in vivo, including the dynamic dynein-LIS1 interaction, as well as the coordination between cargo binding and dynein activation. The mechanisms of dynein regulation, *e*.*g*., kinesin-mediated localization of dynein-dynactin to the microtubule plus-end [76, 87, 88, 90, 118, 142, 143], the importance of the plus-end dynein-dynactin in cargo transport [47, 76, 88, 89, 91, 92, 144-147], the mechanism of dynein-cargo interaction and cargo transport, are all conserved in principle from fungi to higher eukaryotic cells especially elongated neurons [2, 33-36, 38, 94, 95, 148]. Thus, it will be of interest to test whether the wA-AAA3 and wB-AAA3 mutations cause similar effects in higher eukaryotic cells, especially neurons. It will also be highly worthwhile to examine purified mammalian dynein containing the wA-AAA3 or the wB-AAA3 mutation to further dissect the mechanism of the AAA3-involved dynein regulation in vitro in the context of other known factors involved in dynein activation.

## Materials and Methods

### *A. nidulans* strains and media

*A. nidulans* strains used in this study are listed in Table S1. Solid rich medium was made of either YAG (0.5% yeast extract and 2% glucose with 2% agar) or YAG+UU (YAG plus 0.12% uridine and 0.12% uracial). Genetic crosses were done by standard methods. Solid minimal medium containing 1% glucose was used for selecting progeny from a cross. For live cells imaging experiments, cells were cultured in liquid minimal medium containing 1% glycerol for overnight at 32°C. Benomyl was used at the final concentration of 2.4 μg/ml, and the solvent for benomyl (95% EtOH) was used as a control. All the biochemical analyses were done using cells grown at 32°C for overnight in liquid YG rich medium (0.5% yeast extract and 2% glucose). For a few experiments using strains containing the *alcA-nudK*^Arp1^ (Arp1 conditional null) allele, we harvest spores from the solid minimal medium containing 1% glycerol and cultured them in liquid minimal medium containing 1% glucose for imaging analysis.

### Live cell imaging and analyses

All images were captured using an Olympus IX73 inverted fluorescence microscope linked to a PCO/Cooke Corporation Sensicam QE cooled CCD camera. An UPlanSApo 100x objective lens (oil) with a 1.40 numerical aperture was used. A filter wheel system with GFP/mCherry-ET Sputtered series with high transmission (Biovision Technologies) was used. For all images, cells were grown in the LabTek Chambered #1.0 borosilicate coverglass system (Nalge Nunc International, Rochester, NY). The IPLab software was used for image acquisition and analysis. Image labeling was done using Microsoft PowerPoint and/or Adobe Photoshop. For quantitation of signal intensity, a region of interest (ROI) was selected and the Max/Min tool of the IPLab program was used to measure the maximal intensity within the ROI. The ROI box was then dragged to a nearby area inside the cell to take the background value, which was then subtracted from the intensity value. Hyphae were chosen randomly from images acquired under the same experimental conditions. For measuring the signal intensity of a MT plus-end comet formed by GFP-dynein or NudF/LIS1-GFP proteins, only the comet closest to hyphal tip was measured. For measuring GFP-dynein signal intensity at septa, usually only the septum most proximal to the hyphal tip was measured. Images were taken at room temperature immediately after the cells were taken out of the incubators. Cells were cultured overnight in minimal medium with 1% glycerol and supplements at 32°C or 37°C (all experiments using *nudF*6 strains and controls were done at 37°C). Note that the *nudF*6 mutant is temperature sensitive; it forms a tiny colony lacking asexual spores at a higher temperature (typical of a *nud* mutant), but some spores are produced at its semipermissive temperature of 32°C. Thus, for experiments involving *nudF*6, we harvested spores at 32°C and cultured them at 37°C for imaging analysis.

### Constructing the wA-AAA3 (*nudA*^K2599A^) and wB-AAA3 (*nudA*^E2663Q^) mutants and other strains containing the mutant alleles

For making the AAA3 Walker A (wA-AAA3) mutant *nudA*^K2599A^, we used fusion PCR to make a DNA fragment containing the K2599A mutation with the following primers: WAF (5’-TTCTGGTGCAACCATGACACTGTTTGCCG-3’), WAR (5’-GTGTCATGGTTGC ACCAGAACCGGGAGGAC-3’), NudA58[100] and NdA310 [100]. For making the AAA3 Walker B (wB-AAA3) mutant *nudA*^E2663Q^, we used fusion PCR to make a DNA fragment containing the E2663Q mutation with the following primers: WBF (5’-ATCTTCTGCGATCAAATCAACCTGCCGGCTC-3’), WBR (5’-CAGGTTGATTTG ATCGCAGAAGATAACCAGCCAA-3’), NudA58 [100] and NdA310 [100]. Each fragment was co-transformed with a selective marker *pyrG* fragment into the RQ2 strain containing GFP-dynein HC (NudA), mCherry-RabA and the Ku70 deletion allele Δ*nkuA*-*argB*, which facilitates the selection of transformants in which transformed DNA fragments underwent homologous integration into the genome. Several transformants with growth defect were selected and the mutations confirmed by sequencing analysis of the genomic DNA fragments. For making the strains containing p150-GFP and the wA-AAA3 or the wB-AAA3 allele, the fusion PCR product containing the wA-AAA3 or the wB-AAA3 mutation was co-transformed with the p150-GFP-*AfpyrG* fragment [33] into the XY42 strain containing mCherry-RabA and Δ*nkuA*-*argB*. Transformants with growth defect were selected and the mutations confirmed by sequencing analysis. For making the strains containing both the dynein heavy chain tail mutation *nudA*^F208V^ and the wA-AAA3 or the wB-AAA3 allele, the fusion PCR product containing the wA-AAA3 or the wB-AAA3 mutation was co-transformed with the *A. nidulans pyrG* fragment into the RQ338 strain containing GFP-*nudA*^F208V^, mCherry-RabA and Δ*nkuA*-*argB*. Transformants with growth defect were selected and the mutations confirmed by sequencing analysis.

Strains containing multiple mutant alleles were constructed by genetic crosses. Genetic crosses were done by standard methods, and progeny with desired genotypes were selected based on colony phenotype, imaging analysis, Western analysis, diagnostic PCR, and/or sequencing of specific regions of the genomic DNA.

### In vitro dynein motility assay using *A. nidulans* extract

About 0.25 g of hyphal mass was harvested from overnight culture for each sample, and cell extract was prepared using 150 μl of lysis buffer (30 mM HEPES pH 7.3, 50 mM potassium acetate, 2 mM magnesium acetate, 1 mM EGTA, 10% glycerol, 1 mM DTT, and 0.2 mM ATP). Cell extracts were then centrifuged at 20,000 *g* for 30min at 4°C to remove cell debris, and supernatant was kept on ice for motility assays, or snap-frozen with liquid nitrogen and stored at -80°C if motility assays cannot be performed on the same day. Immediately prior to imaging, the cell lysates were warmed to room temperature and diluted 10 times in the lysis buffer containing 1 mM ATP, 1 mM Trolox, 20 μM Taxol, 10 mM DTT, 1 mg/mL BSA, 1 mg/mL β-casein, 15 μg/mL catalase, 40 μM glucose, and 40 μg/mL glucose oxidase.

To make X-rhodamine labeled microtubule, un-labeled tubulin (Cytoskeleton, Inc) and X-rhodamine-labeled tubulin (Cytoskeleton, Inc) are mixed with a ratio of ∼12:1 in BRB80 buffer (80 mM PIPES, pH 7.0, 1 mM EGTA, and 1 mM MgCl_2_) with 2 mM GTP. After incubating for 12 min at 35°C, the polymerized microtubule was stabilized by adding 20 μM taxol and stored at room temperature.

To prepare flow chambers, 75 mm x 25 mm glass microscope slides and 18 mm x 18 mm coverslips with #1.5 thickness were treated as described [149]. In summary, both the microscope slides and coverslips were pre-cleaned with alkaline in bath sonicator followed by washing with ultrapure water and treating in plasma clear with RF coil set on “high” for 5 min. The pre-cleaned coverslips were further aminosilanized with 2% APTES (Sigma-Aldrich). To assemble the slide chamber, a set of parafilm strips were inserted between the aminosilane surface-coated coverslip and pre-cleaned slide to generate an approximate volume of 15-20 μL and the chamber was then formed after being heat sealed. The aminosilanized surface of the slide chamber was further activated with 8% glutaraldehyde (Sigma-Aldrich) to enhance microtubule binding prior to an experiment.

To image dynein motility on microtubule, 20 μL of ∼0.1 mg/mL microtubule was flushed into the glutaraldehyde treated chamber from one side and incubate at room temperature for 90 seconds, 20 μL BRB80 with 20 μM taxol was then added from the same side and the chamber was incubated at room temperature for 20 min to allow microtubule binding. The flow chamber was then blocked with 2 mg/mL β-casein and 20 μM Taxol for 5 min, and the cell lysates were added in the presence of either 1 mM or 3 mM ATP. Time lapse movies were acquired immediately with either an Olympus IX73 inverted fluorescence microscope linked to a PCO/Cooke Corporation Sensicam QE cooled CCD camera or an Olympus IX83 microscope equipped with a one-line motorized IX3 cellTIRF illuminator and quad laser TIRF set, outfitted with an ORCA flash 4.0 v3 sCMOS camera (Hamamatsu), a 100x 1.50-NA objective, and CellSens software. A 488-nm laser and a 561-nm laser were used for acquiring images of GFP-labeled dynein and X-Rhodamine-labeled microtubules respectively. Several 1.5 to 2 min movies were acquired at room temperature per chamber with 1 second intervals of each frame. Velocities were measured by using kymographs generated by the Fiji/ImageJ software (the National institutes of Health).

### Statistical analysis

All statistical analyses were done using GraphPad Prism 8 for Mac (version 8.0.0, 2018). The D’Agostino & Pearson normality test was performed on all data sets. For all data sets, non-parametric tests were used without assuming Gaussian distribution. Specifically, the Kruskal-Wallis ANOVA test (unpaired) with Dunn’s multiple comparisons test was used for analyzing multiple data sets. The Mann-Whitney test (unpaired, two tailed) was used for analyzing the two data sets. Note that adjusted p values were generated from the Kruskal-Wallis ANOVA test with Dunn’s multiple comparisons test. In all figures, **** indicates p<0.0001, *** indicates p<0.001, ** indicates p<0.01, * indicates p<0.05 *, and “ns” indicates p>0.05.

## Acknowledgements

We thank Drs. Berl Oakley, Stephen Osmani and Miguel Peñalva for Aspergillus strains. We would also like to thank Dr. Peter Höök (former postdoctoral fellow from Dr. Richard Vallee’s lab) and Dr. Samara Reck-Peterson for informing us many years ago that *nudA*^R3086^ represents an arginine-finger. This work was funded by the National Institutes of Health RO1 GM121850 (to X. Xiang) and RO1 GM134104 (to J. Rotty). J. Rotty was also supported by Uniformed Services University start-up fund. Disclaimer: The opinions and assertions expressed herein are those of the author(s) and do not necessarily reflect the official policy or position of the Uniformed Services University or the Department of Defense.

## Author contributions

R.Q. and X.X. conceived the experiments. R.Q., J.Z., J.D.R. and X.X. carried out the experiments. X.X. and R.Q. wrote the manuscript, which was read and edited by all authors. All authors helped shape the research, analysis and manuscript.

## Competing interests

The authors declare no competing interest.

## Online Supplemental Materials

**Table S1.** *Aspergillus nidulans* strains used in this study.

| Strain | Genotype | Source |
| --- | --- | --- |
| JZ383 | $\Delta nudF::pyr4$; GFP-*nudA*<sup>HC</sup>; *argB2::[argB\*-alcAp::mCherry-RabA]* | [68] |
| LZ36 | GFP-*nudA*<sup>R3086C</sup>; $\Delta nkuA::argB$; *pyroA4* | [100] |
| RQ2 | GFP-*nudA*<sup>HC</sup>; *argB2::[argB\*-alcAp::mCherry-RabA]*; $\Delta nkuA::argB$; *pyrG89*; *pyroA4*; *yA2* | [109] |
| RQ8 | GFP-*nudA*<sup>F208V</sup>; *argB2::[argB\*-alcAp::mCherry-RabA]*; *pyrG89*; *wA2* | [109] |
| RQ165 | GFP-*nudF-AfpYrG*; *argB2::[argB\*-alcAp::mCherry-RabA]*; $\Delta nkuA::argB$; *pyroA4*; *pyrG89*; *wA2* | [68] |
| RQ275 | *nudF6*; *gpdA-ΔC-hookA-S-AfpYrG*; GFP-*nudA*<sup>HC</sup>; *pyrG89*; *yA2*; possibly $\Delta nkuA::argB$ | [68] |
| RQ287 | GFP-*nudA*<sup>HC</sup>; *gpdA-ΔC-hookA-S-AfpYrG*; *yA2* | [68] |
| XX60 | $\Delta nudA::pyrG$; *pyrG89* | [105] |
| XX222 | GFP-*nudA*<sup>HC</sup>; *argB2::[argB\*-alcAp::mCherry-RabA]*; *yA2*; *pantoB100* | [89] |
| XX514 | GFP-*nudA*<sup>HC</sup>; $\Delta yA::NLS-DsRed$; possibly $\Delta nkuA::argB$ | [68] |
| XX634 | $\Delta nudF::pyr4$; GFP-*nudA*<sup>R1602E, K1645E</sup>; *argB2::[argB\*-alcAp::mCherry-RabA]*; possibly $\Delta nkuA::argB$ | [68] |
| XY13 | p150-GFP-*AfpYrG*; $\Delta nkuA::argB$; *pyrG89*; *pyroA4* | [90] |
| XY42 | *argB2::[argB\*-alcAp::mCherry-RabA]*; $\Delta nkuA::argB$; *pyrG89*; *yA2*; *pantoB100* | [150] |
| JZ504 | GFP-*nudA*<sup>HC</sup>; $\Delta hookA-AfpYrG$; *argB2::[argB\*-alcAp::mCherry-RabA]*; *pyroA4* | This work |
| RQ102 | GFP-*nudA*<sup>HC</sup>; $\Delta hookA-AfpYrG$; *argB2::[argB\*-alcAp::mCherry-RabA]*; *pantoB100*; *yA2* | This work |
| RQ293 | *nudF6*; *gpdA-ΔC-hookA-S-AfpYrG*; GFP-*nudA*<sup>R3086C</sup> | This work |
| RQ295 | *gpdA-ΔC-hookA-S-AfpYrG*; GFP-*nudA*<sup>R3086C</sup> | This work |
| RQ300 | GFP-*nudA*<sup>K2599A</sup>(*wA-AAA3*); *argB2::[argB\*-alcAp::mCherry-RabA]*; $\Delta nkuA::argB$; *pyroA4*; *yA2* | This work |
| RQ301 | GFP-*nudA*<sup>E2663Q</sup>(*wB-AAA3*); *argB2::[argB\*-alcAp::mCherry-RabA]*; $\Delta nkuA::argB$; *pyroA4*; *yA2* | This work |
| RQ306 | $\Delta nudF::pyr4$; GFP-*nudA*<sup>K2599A</sup>(*wA-AAA3*); *argB2::[argB\*-alcAp::mCherry-RabA]* | This work |
| RQ307 | $\Delta nudF::pyr4$; GFP-*nudA*<sup>E2663Q</sup>(*wB-AAA3*); *argB2::[argB\*-alcAp::mCherry-RabA]* | This work |
| RQ309 | GFP-*nudA*<sup>K2599A</sup>(*wA-AAA3*); *gpdA-ΔC-hookA-S-AfpYrG* | This work |
| RQ311 | GFP-*nudA*<sup>K2599A</sup>(*wA-AAA3*); $\Delta yA::NLS-DsRed$; possibly $\Delta nkuA::argB$ | This work |
| RQ312 | GFP-*nudA*<sup>E2663Q</sup>(*wB-AAA3*); *gpdA-ΔC-hookA-S-AfpYrG* | This work |
| RQ314 | GFP-*nudA*<sup>E2663Q</sup>(*wB-AAA3*); $\Delta yA::NLS-DsRed$; possibly $\Delta nkuA::argB$ | This work |
| RQ317 | GFP-*nudA*<sup>K2599A</sup>(*wA-AAA3*); *gpdA-ΔC-hookA-S-AfpYrG*; *argB2::[argB\*-alcAp::mCherry-RabA]* | This work |
| RQ318 | GFP-*nudA*<sup>K2599A</sup>(*wA-AAA3*); *gpdA-ΔC-hookA-S-AfpYrG*; *nudF6* | This work |
| RQ323 | *nudA*<sup>K2599A</sup>(*wA-AAA3*); p150-GFP-*AfpYrG*; $\Delta nkuA::argB$; *argB2::[argB\*-alcAp::mCherry-RabA]*; *pantoB100*; *yA2* | This work |
| RQ324 | *nudA*<sup>E2663Q</sup>(*wB-AAA3*); p150-GFP-*AfpYrG*; $\Delta nkuA::argB$; *argB2::[argB\*-alcAp::mCherry-RabA]*; *pantoB100*; *yA2* | This work |
| RQ328 | GFP-*nudA*<sup>E2663Q</sup>(wB-AAA3); *argB2::[argB\*-alcAp::mCherry-RabA]*; *ΔhookA-AfpYrG* | This work |
| RQ329 | GFP-*nudA*<sup>K2599A</sup>(wA-AAA3); *argB2::[argB\*-alcAp::mCherry-RabA]*; *ΔhookA-AfpYrG* | This work |
| RQ330 | GFP-*nudA*<sup>K2599A</sup>(wA-AAA3); *argB2::[argB\*-alcAp::mCherry-RabA]* | This work |
| RQ331 | GFP-*nudA*<sup>E2663Q</sup>(wB-AAA3); *argB2::[argB\*-alcAp::mCherry-RabA]* | This work |
| RQ332 | GFP-*nudA*<sup>K2599A</sup>(wA-AAA3); *alcAp-nudK<sup>Arp1</sup>-pyr4*; possibly *argB2::[argB\*-alcAp::mCherry-RabA]*; possibly *p25-S* | This work |
| RQ333 | GFP-*nudA*<sup>E2663Q</sup>(wB-AAA3); *alcAp-nudK<sup>Arp1</sup>-pyr4*; possibly *argB2::[argB\*-alcAp::mCherry-RabA]*; possibly *p25-S* | This work |
| RQ338 | GFP-*nudA*<sup>F208V</sup>; *argB2::[argB\*-alcAp::mCherry-RabA]*; *ΔnkuA::argB*; *pyrG89*; *pyroA4* | This work |
| RQ343 | GFP-*nudA*<sup>K2599A, F208V</sup>; *argB2::[argB\*-alcAp::mCherry-RabA]*; *ΔnkuA::argB*; *pyroA4* | This work |
| RQ344 | GFP-*nudA*<sup>E2663Q, F208V</sup>; *argB2::[argB\*-alcAp::mCherry-RabA]*; *ΔnkuA::argB*; *pyroA4* | This work |
| RQ354 | GFP-*nudF-AfpYrG*; *nudA*<sup>K2599A</sup>(wA-AAA3); *argB2::[argB\*-alcAp::mCherry-RabA]*; possibly *ΔnkuA::argB*; *pyrG89* | This work |
| RQ355 | GFP-*nudF-AfpYrG*; *nudA*<sup>E2663Q</sup>(wB-AAA3); *argB2::[argB\*-alcAp::mCherry-RabA]*; possibly *ΔnkuA::argB*; *pyrG89* | This work |
| XX632 | GFP-*nudA*<sup>R3086C</sup>; *ΔnudF::pyr4*; *argB2::[argB\*-alcAp::mCherry-RabA]*; possibly *ΔnkuA::argB*; possibly *pyrG89* | This work |
| XX644 | GFP-*nudA*<sup>K2599A</sup>(wA-AAA3); *nudF6*; *argB2::[argB\*-alcAp::mCherry-RabA]*; possibly *ΔnkuA::argB* | This work |
| XX684 | GFP-*nudF-AfpYrG*; *nudA*<sup>K2599A</sup>(wA-AAA3); *gpdA-ΔC-hookA-S*; possibly *ΔnkuA::argB*; possibly *pyrG89*; possibly *argB2::[argB\*-alcAp::mCherry-RabA]* | This work |
| XX689 | GFP-*nudF-AfpYrG*; *nudA*<sup>E2663Q</sup>(wB-AAA3); *gpdA-ΔC-hookA-S*; possibly *ΔnkuA::argB*; possibly *pyrG89*; possibly *argB2::[argB\*-alcAp::mCherry-RabA]* | This work |

**Figure S1.**
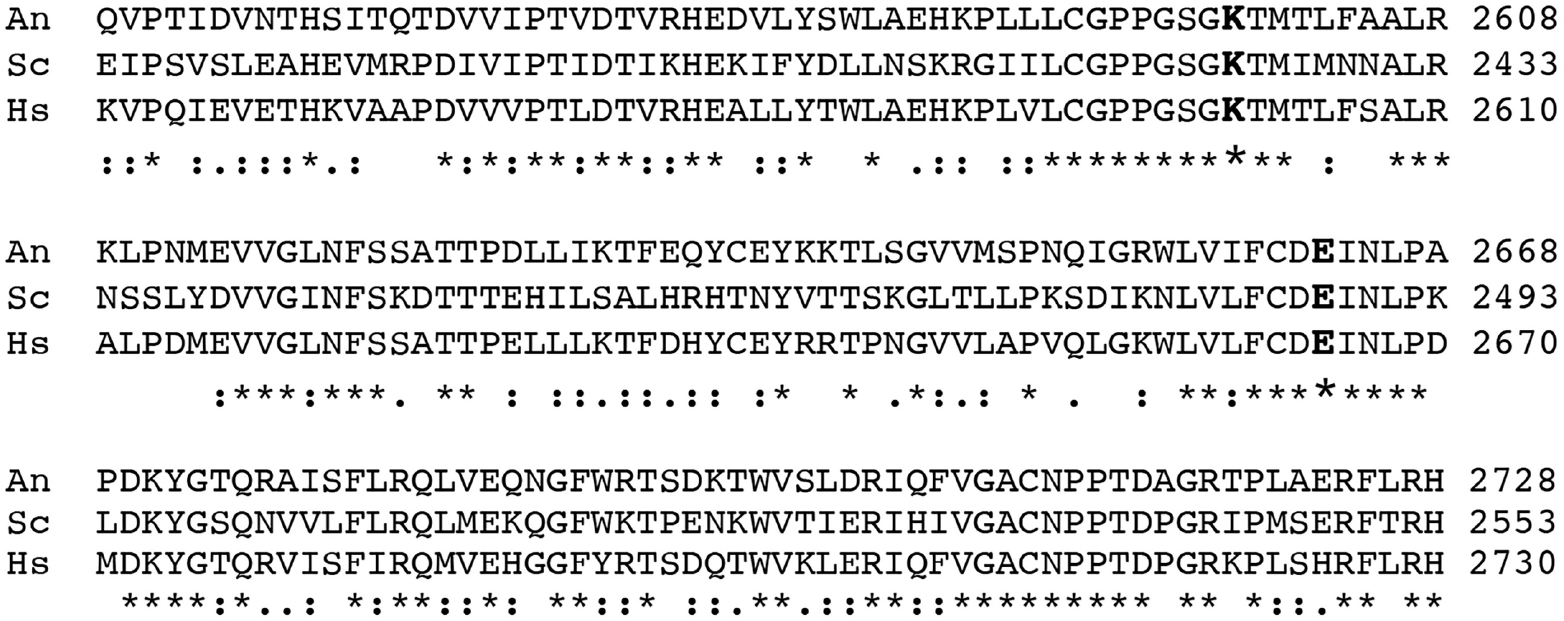
A part of the sequence alignment of dynein heavy chains from *Aspergillus nidulans* (An), the budding yeast or *Saccharomyces cerevisiae* (Sc) and human or Homo sapiens (Hs). The alignment was done using Clustal Omega (https://www.ebi.ac.uk/Tools/msa/clustalo/). Residues that are identical (*), strongly similar (:) or weakly similar (.) are indicated. The Lysine (K) of AAA3 Walker A and the Glutamic acid (E) of AAA3 Walker B are shown using bold characters.

**Figure S2.**
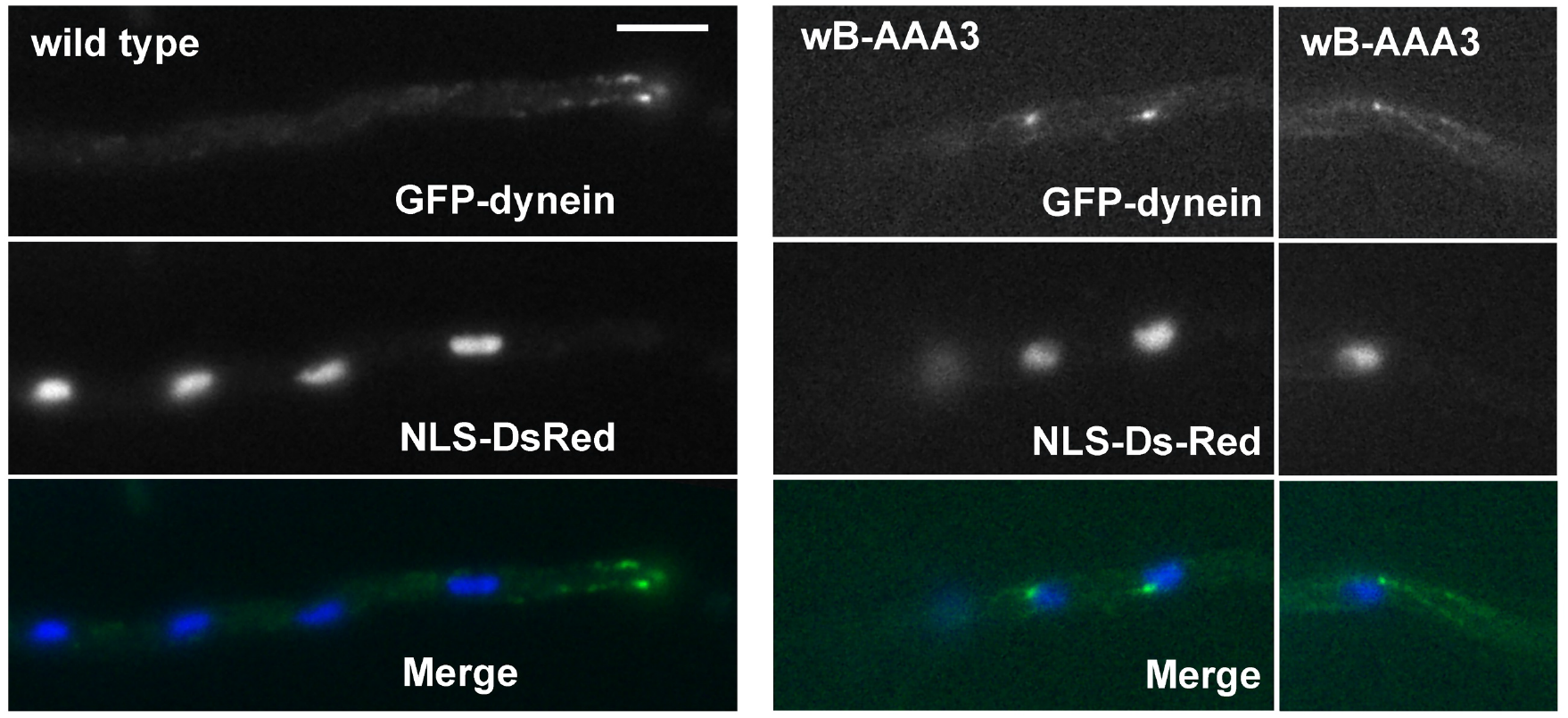
The spindle-pole body (SPB) localization of the wB-AAA3 dynein on nuclei. The wB-AAA3 dynein is labeled with GFP. Nuclei are labeled with NLS-DsRed and pseudo-colored in blue. A wild-type hypha is shown as a control, in which GFP-dynein mainly forms microtubule plus-end comets. Bar, 5 μm.

**Figure S3.**
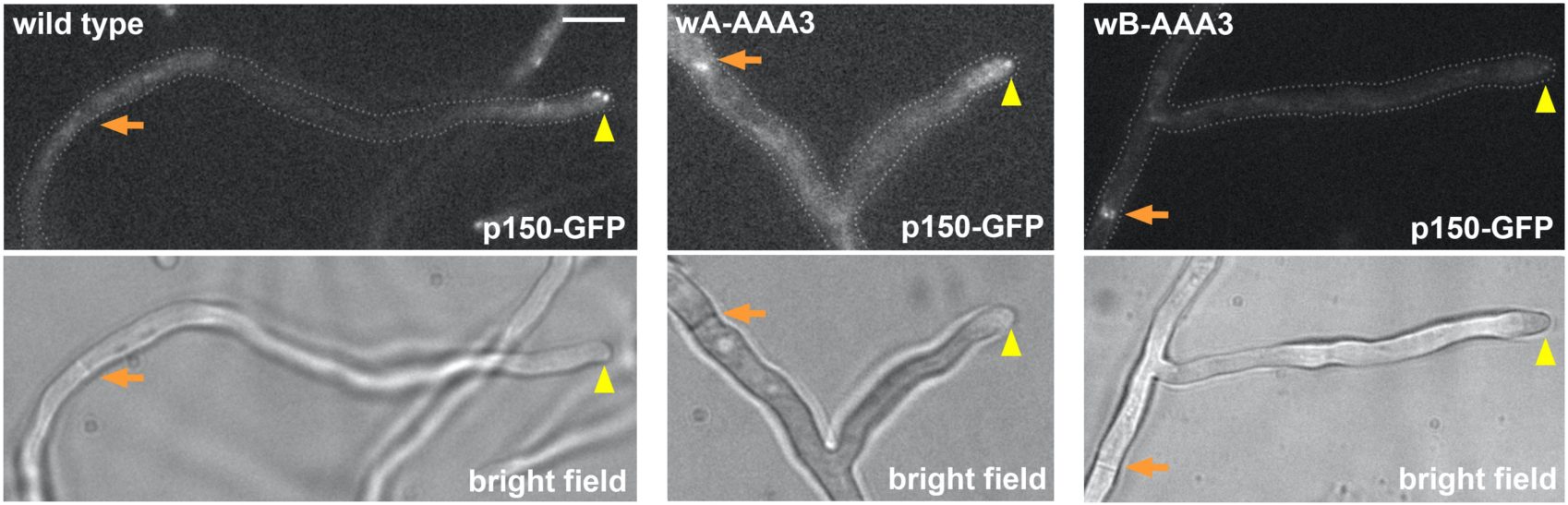
Localization of dynactin p150-GFP in wild type, the wA-AAA3 mutant and the wB-AAA3 mutant. Bright-field images are shown below to indicate hyphal shape and position of septum. Hyphal tip is indicated by a yellow arrowhead and septum by a light brown arrow. Bar, 5 μm.

**Figure S4.**
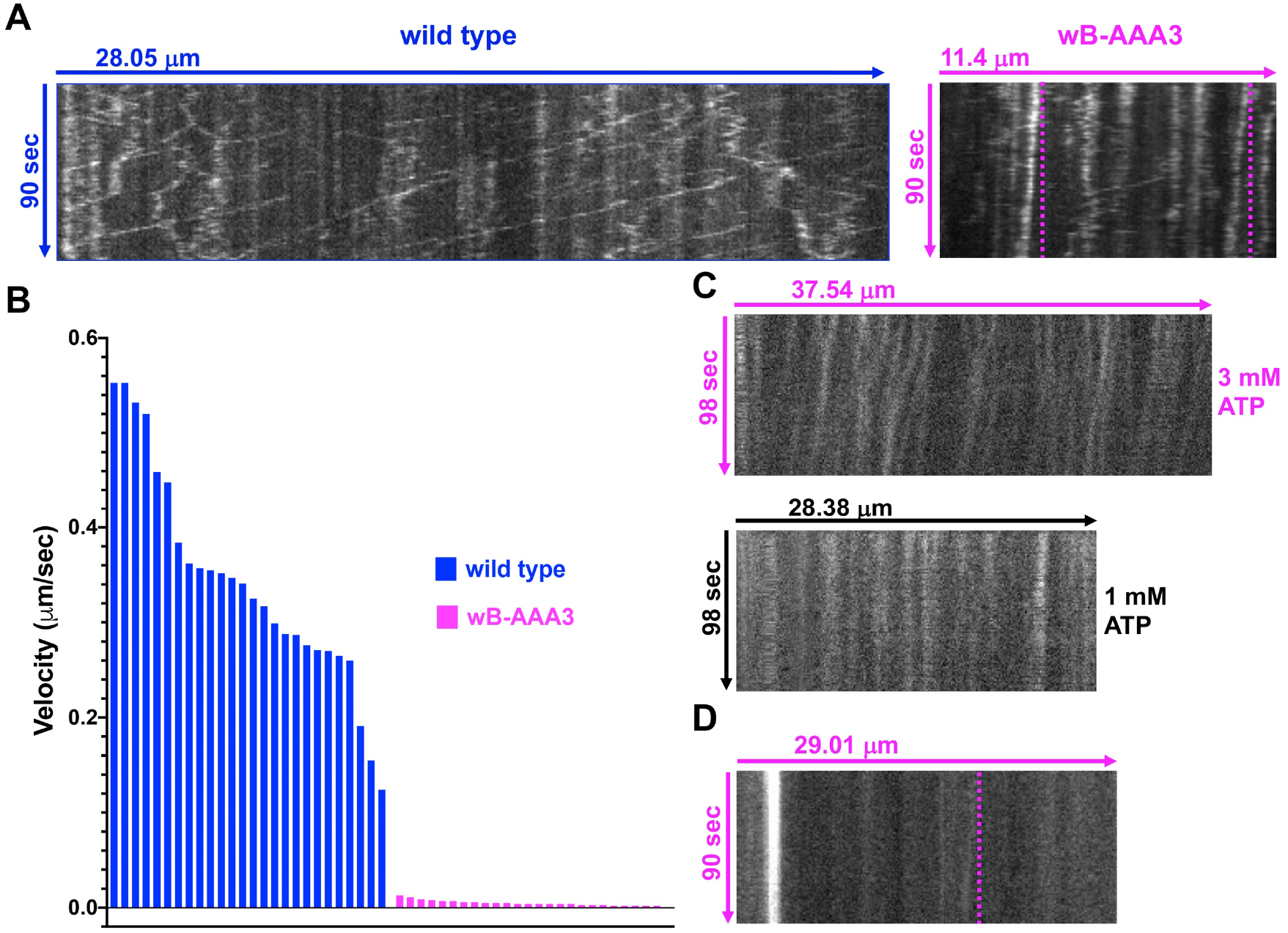
The wB-AAA3 mutation dramatically decreases the speed of dynein motility in vitro. (A) kymographs showing the in vitro movements of wild-type dynein and wB-AAA3 dynein. For wild-type dynein, we used cell extract from a strain containing the *gpdA*-ΔC-*hookA*-S fusion to ensure that GFP-dynein molecules predominantly move toward the minus ends. For wB-AAA3 dynein, we did not include the *gpdA*-ΔC-*hookA*-S fusion because the wB-AAA3 dynein accumulates at the minus ends even in the absence of HookA, and the wB-AAA3, *gpdA*-ΔC-*hookA*-S double mutant is very sick. (B) Velocity of wild-type dynein verses that of the wB-AAA3 dynein (n=26 for wild-type dynein; n=25 for the wB-AAA3 mutant dynein). Note that the wB-AAA3 dynein moves with an extremely low speed. The mean ranks of these two sets of data are significantly different (p<0.0001, unpaired, Mann-Whitney test, Prism 8). (C) Kymographs of the wB-AAA3 dynein movements in 3 mM ATP or 1 mM ATP. All the velocity data from the wB-AAA3 dynein were obtained in the in vitro motility assays with 3 mM ATP, although 1 mM ATP was used for the wild-type dynein. This is because the wB-AAA3 dynein was not motile if 1 mM ATP is used. (D) A kymograph showing the occasional slow movement of a p150-GFP signal toward a septum in vivo. The very bright line corresponds to the position of the septum (note that the wB-AAA3 mutation causes both dynein and dynactin to accumulate at septa).

**Figure S5.**
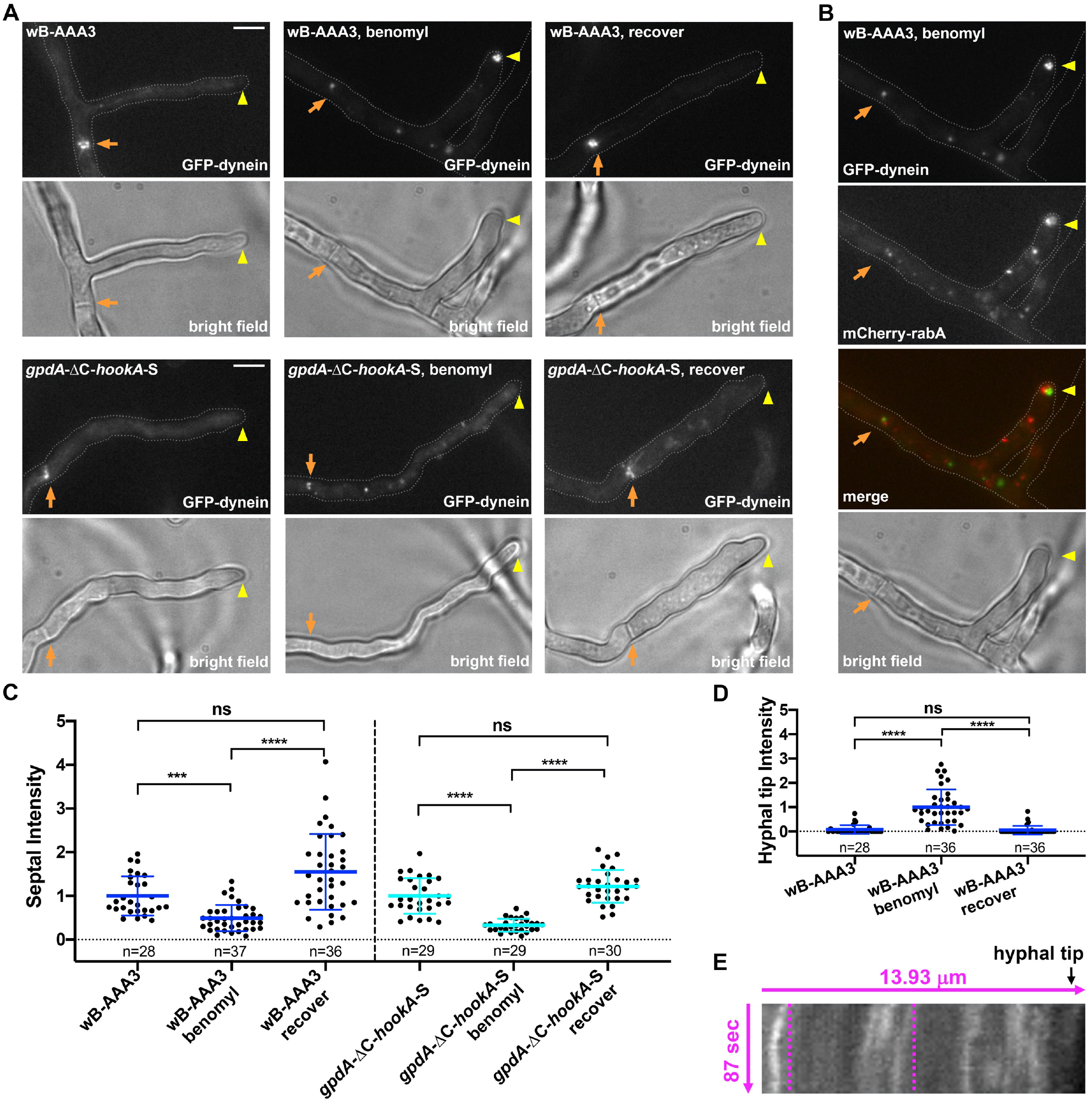
Effect of benomyl on the localization of wB-AAA3 dynein. (A) images of the wB-AAA3 dynein 3 hours after benomyl treatment and 4 hours after recovery. The *gpdA*-ΔC-*hookA*-S strain containing wild-type dynein was used as a control (note that dynein is driven to septa upon overexpression of ΔC-HookA in the *gpdA*-ΔC-*hookA*-S strain). Although not indicated specifically, both the wB-AAA3 and *gpdA*-ΔC-*hookA*-S samples were treated with EtOH (the solvent of benomyl) as controls. Bright-field images are shown below to indicate hyphal shape and position of septum. Hyphal tip is indicated by a yellow arrowhead and septum by a light brown arrow. Bar, 5 μm. (B) Images showing the partial overlap of the wB-AAA3 dynein signals and the mCherry-RabA signals benomyl treatment. (C)(D) Quantitative analyses on the septal and hyphal-tip dynein intensity. In (C), the average values for dynein without benomyl treatment are set as 1. In (D), the average values for dynein in benomyl-treated cells are set as 1. Scatter plots with mean and S.D. values were generated by Prism 8. ****p<0.0001, ***p<0.001 (Kruskal-Wallis ANOVA test with Dunn’s multiple comparisons test, unpaired). (E) A kymograph showing a few wB-AAA3 dynein signals moving away from the hyphal tip during recovery from the benomyl treatment.

## Notes

### Competing Interest Statement

The authors have declared no competing interest.

